# Interaction of light-dark and dietary cues prime *Drosophila* ovarian stem cell properties by precisely tuning Drp1 recruitment and organization on mitochondria

**DOI:** 10.64898/2026.09.04.749535

**Authors:** Yoshita Sriramkumar, Rhea Wali, Kasturi Mitra

## Abstract

The quiescent adult stem cells are activated to maintain tissue integrity. Mitochondria have emerged as critical regulators of stem/progenitor cells. Scattered literature suggests existence of a high-potency stemness state with characteristic mitochondrial structure-function properties that may ‘prime’ stem cell activation. Precise tuning of mitochondrial structure by the mitochondrial fission protein, Drp1, can prime a stem cell state for neoplasticity in vitro. However, the in vivo significance and regulation of such a proposed ‘primed’ stemness state remains elusive. We use quantitative genetic and cell biological approaches to demonstrate that light-dark cycling driven precise tuning of Drp1 primes the *Drosophila* ovarian germline stem cell (GSC) response to protein-rich dietary supplement towards stimulating egg production. This can be reversed by repressing mitochondrial fusion protein, Opa1, and is disrupted in the absence of functional Cry protein that maintains light-dark rhythm. Mechanistically, this is achieved by light or dark allowing protein-rich supplements to stimulate distinct characteristic recruitment and organization of Drp1 on mitochondria in GSC subpopulations. Screening of essential amino acids reveals the importance of interaction between Threonine supplementation, Cry, Drp1 and Opa1 in such priming of the GSCs and egg production. The priming of the ovarian follicle stem cells happens at a different tuned level of Drp1. Such Drp1-tuning driven priming of *Drosophila* ovarian stem cells reverses the inhibitory impact of diabetes-like condition on egg development in the presence of protein-rich supplement. Thus, our study revealing the paradigm of adult stem cell regulation by integration of light-dark and dietary cues via precise mitochondrial modulation, is significant in regenerative health and degenerative diseases.

## Introduction

The small number of adult stem cells residing at their respective niche are largely quiescent. They are activated to self-renew, proliferate and differentiate on demand for maintaining tissue integrity and repairing damage (Fuchs and Blau 2020). Various tissues undergo adult stem cell exhaustion or accumulate dysfunctional stem cells during aging (Rando et al. 2025; Liu et al. 2022; Ishibashi et al. 2020). The tissue resident adult stem cells can undergo neoplastic transformation under oncogenic insults, which differentiate into bulk tumor cells to form tumors (Brian Spurlock, Parker, et al. 2021). A tissue may harbor multiple lineage committed stem cell types, while the stem cell state is regulated by niches and environmental milieu (Merrell and Stanger 2016; Poss and Tanaka 2024).

Mitochondria are homeostatic organelles that can integrate internal and external cellular cues with cellular signaling pathways and thus contribute to cell fate determination (Suomalainen and Nunnari 2024). Mitochondrial structure and function impact each other as regulated by a set of molecules that either cause mitochondria to undergo fission or fusion. The action of the primary driver of mitochondrial fission, the dynamin related protein 1 (Drp1), is modulated by specific regulators, and is opposed by mitochondrial fusion primarily regulated by Opa1 and Mitofusins (Quintana-Cabrera and Scorrano 2023). While the mitochondrial fusion regulators are bonafide mitochondrial proteins, Drp1 needs to be recruited to the mitochondrial surface by bonafide mitochondrial proteins, MFF and Fis-1, to render their mitochondrial fission activity (Tábara et al. 2025; Giacomello et al. 2020). A large body of literature shows that specific mitochondrial structure and function are critical for stem/progenitor cell regulation (Chakrabarty and Chandel 2021; Folmes and Terzic 2016). Alteration of mitochondrial fission and fusion molecules modulates self-renewal and differentiation of stem/progenitor cells of various lineages (Lisowski et al. 2018; Khacho and Slack 2017; Verma et al. 2026).

Drp1 activity / recruitment has been shown to maintain a certain metabolic state which is associated with a proposed ‘primed’ state of hematopoietic stem cells that is poised for activation and proliferation in mice (Liang et al. 2020; Qiu and Ghaffari 2022; Qiu et al. 2021). Such a priming can be considered to underlie the boost in neural stem cell functionality and neurogenesis driven by Drp1 knockdown in mice (Iwata et al. 2020). However, Drp1 gene ablation can be deleterious to stem cell properties across lineages (Khacho et al. 2016; Hinge et al. 2020; Cieslar-Pobuda and Caglayan 2025). We reported that maintaining a tuned level of Drp1 substantially elevates stimulated self-renewal and proliferation of stem cells in skin lineage in vitro (Spurlock et al. 2021b). Such tuned levels of Drp1 is required to maintain a characteristic mitochondrial form to specify stemness by modulating gene-expression ( Spurlock et al. 2021b; Saini et al. 2024). Therefore, mitochondrial priming of stemness, by maintaining mitochondrial fission at a level that can be activated as well as repressed as needed, can potentially make the adult stem cells more potent. The mitochondrial primed state of stemness appears to be modulated by mitochondrial potential in mouse and human, and can be potentially isolated based on mitochondrial potential (Sukumar et al. 2016; Spurlock et al. 2021a; Spurlock et al. 2019). Therefore, the reports of linking mitochondrial content and membrane potential linking to stemness in various lineages across various organisms could also reflect existence of a mitochondrial primed state (Mohamed Haroon et al. 2021; Pan et al. 2024; Mansell et al. 2021; Papa et al. 2020). Mitochondrial transmembrane potential is maintained by utilizing dietary carbon sources. Although metabolic regulation of stem cells is widely being studied across various lineages (Jackson and Finley 2024), the carbon source requirement from the diet to support the transmembrane potential of mitochondrial primed state remains unknown.

Nutrient availability can impact mitochondrial fission-fusion and nutrient utilization can be regulated by modulating the mitochondrial fission-fusion balance (Schrepfer and Scorrano 2016; Liesa and Shirihai 2013; Rambold et al. 2011; Xia et al. 2024). However, this relationship remains tissue and context-specific without general consensus, likely because mitochondria is wired differently in different tissues. Dietary metabolism is under the influence of circadian and diurnal rhythm through syncing of the fed-fast and light-dark rhythm with central and peripheral tissues exhibiting distinct responses to changes of the environment (Reinke and Asher 2019; Patke et al. 2020; Bass 2024). Disruption of light-dark rhythms by genetic alteration of core-circadian regulators like Clock and Bmal, impact Drp1 functionality in various tissue types (Schmitt et al. 2018; Xu et al. 2022; Jacobi et al. 2015). Also, the light-sensing molecule, Cryptochrome, has been shown to regulate metabolism apart from their conventional circadian roles (Lamia et al. 2009; Zhang et al. 2010). Recent evidence suggests Cry has a clock-independent role in governing the metabolic state in pluripotent stem cells and their differentiated counterparts (Dierickx et al. 2017; Sato et al. 2023). However, any link between Cry and Drp1 remains to be established.

Tissue resident adult stem/progenitor cells have been widely studied in blood, brain, skin, skeletal muscle, intestines, lung and ovaries. Direct evidence of *in vivo* existence of the mitochondrial primed stem cell state regulated by tuning Drp1 is still missing. To fill this gap, we chose the simple and well-defined model system of *Drosophila* ovary with only two kinds of stem cells, germline stem cells (GSCs) and somatic follicle stem cells (FSCs). The GSCs and FSCs have been studied extensively to serve as models of adult stem cells, and mitochondrial regulation of both the lineages have been documented (Mitra et al. 2012; Tiwari and Mandal 2021; Uttekar et al. 2024; Tomer et al. 2018). Activation of GSC and FSC lineage lead to egg production that exhibits light-dark rhythm (Manjunatha et al. 2008; Sellix and Menaker 2010; Riva et al. 2026) and can be stimulated by protein-rich diets and attenuated by high sugar diet (Nunes and Drummond-Barbosa 2023; Skorupa et al. 2008; Drummond-Barbosa 2019). However, the molecular interactions of the rhythms of light-dark and diet have not been explored in *Drosophila* ovarian development. Here, we use precise genetic and cell biological approaches to demonstrate that Drp1 tuning to an optimal level sustains a mitochondrial primed state in *Drosophila* ovarian stem cells in an Opa1, Cry and Threonine dependent manner. While the GSCs and FSCs are primed by a different level of Drp1, the GSCs undergo Drp1 tuning in the light or dark cycles to allow specific action of the diet for sustaining their mitosis, self-renewal and differentiation to support egg production.

## Results

### Precise tuning of Drp1 driven mitochondrial regulation primes *Drosophila* ovarian stem cell response to protein-rich dietary supplement towards stimulating egg production

We studied the impact of Drp1 tuning on *Drosophila* ovarian GSC/FSC and egg production by employing quantitative lineage tracing analyses of mutant clones or expressing RNAi. We performed detailed comparative analyses of two Drp1 hypomorphic alleles of different strengths and a functional null allele with the following order of Drp1 activity: wildtype (WT) > *drp1^2^* > *drp1^1^* > *drp1^KG^* (Verstreken et al. 2005; Sandoval et al. 2014). The stem cell lineages were traced using GFP negative homozygous clones created by FLP-FRT strategy where the FRT-40A clone served as wildtype (WT)(See METHODS). The GSCs/FSCs were identified as the first GFP-negative clonal cell in their respective anatomical position in the lineage (Fig S1A). The developmental outcome of the clonal GSCs/FSCs is used to measure their properties (Fig. S1B, details in METHODS) (Spradling 2009; Laws and Drummond-Barbosa 2015). We studied the involvement of protein-rich dietary supplementation by comparing the phenotypes in the presence (stimulated) or absence (basal) of supplemented yeast.

The GSC lineage tracing analyses showed that the basal abundance of *drp1^KG^* mutant clones is lower compared to other drp1-mutants, likely due to clearance of *drp1^KG^* clones without detectable mitochondria (Fig. S1C-D). This defect of the *drp1^KG^*-GSCs is rescued with dietary stimulation (Fig. S1D). Notably, there is an increase in basal loss of all drp1-mutant GSCs, which is reduced with dietary stimulation (Fig. 1A-B). Interestingly, dietary stimulation further increases the already higher basal self-renewal ability of the *drp1^2^* GSCs that have the mildest reduction of Drp1 activity (Fig. 1A-B). This data reveals that the level of Drp1 activity in the *drp1^2^*-GSCs is optimal for self-renewal / proliferation, which when altered reduces self-renewal in WT and other Drp1-mutant GSCs. On the other hand, dietary stimulation maximizes differentiation of the *drp1^1^*- GSCs into their germline progenies and minimizes it in the *drp1^KG^* GSCs, with no basal difference (Fig. 1C). The basal stunted cysts differentiated from the *drp1^2^* or *drp1^1^* GSCs grow normally in stimulated conditions, which fails to happen in the stunted cysts differentiated from *drp1^KG^*-GSCs (Fig. S1E).

**Figure 1.**
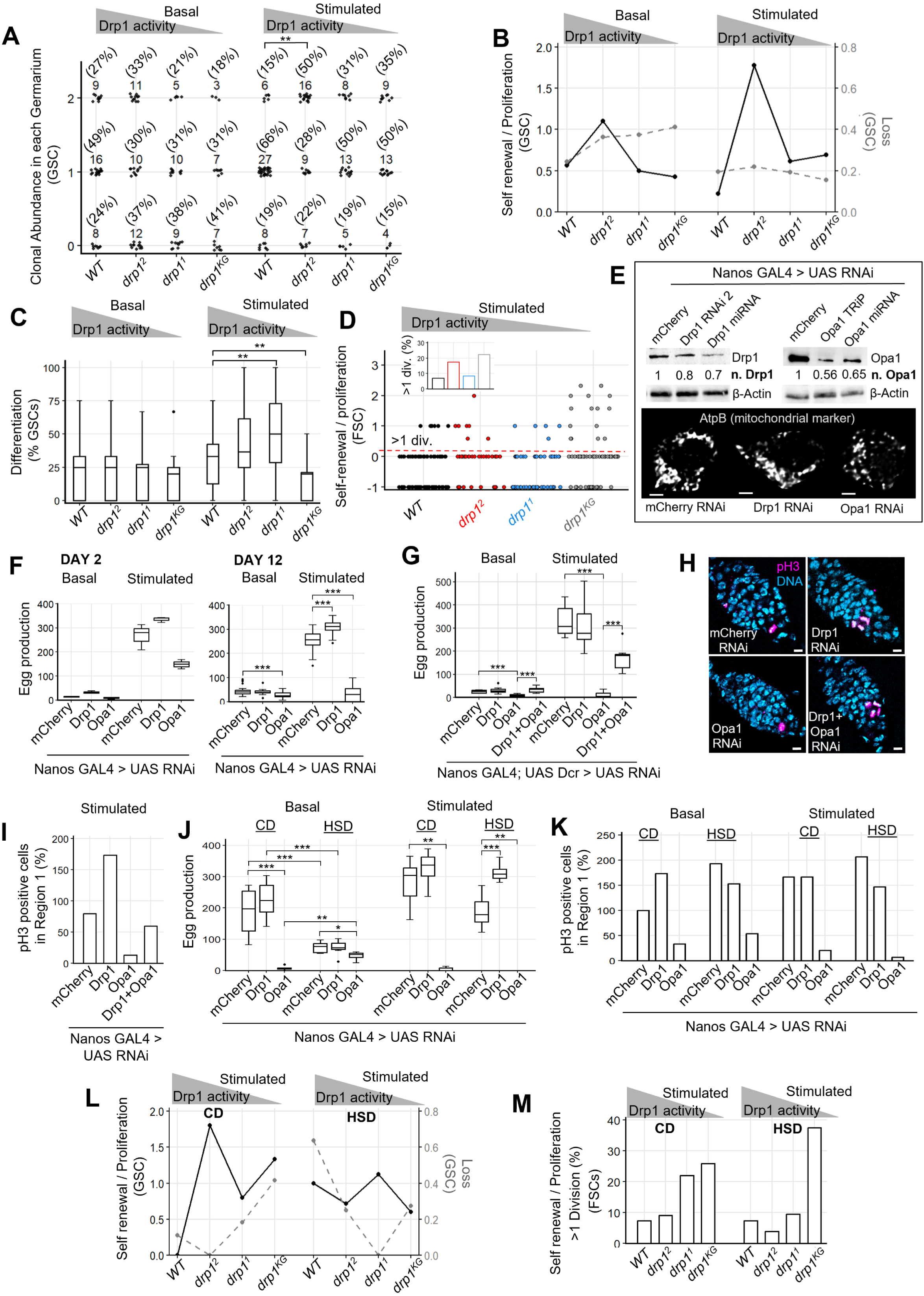
**A**) Dot plots showing clonal GSC abundance in each germarium with 0-, 1-, and 2-GSCs where frequency (%) denoted in parentheses for each genotype; gray triangle depicting decreasing Drp1 activity across the mutants. (N(germaria)-Basal: WT-33, *drp1^2^*-33, *drp1^1^*-24, *drp1^KG^*-17; N(germaria)-Stimulated: WT-41, *drp1^2^*-32, *drp1^1^*-26, *drp1^KG^*-26). **B)** Line plots showing GSC Self-renewal (black) and Loss (grey-dashed), as calculated from A. **C)** Boxplots showing abundance of differentiated clonal cysts (%) in each ovariole for each genotype. (N(ovariole)-Basal: WT-30, *drp1^2^*-29, *drp1^1^*-24, *drp1^KG^*-17; N(ovariole)-Stimulated: WT-40, *drp1^2^*-32, *drp1^1^*-26, *drp1^KG^*-16). **D)** Scatter plot showing FSC Self-renewal / Proliferation for each genotype. Inset shows bar plot of FSCs (%) with >1 division per cyst (N(cyst)-Stimulated : WT-86, *drp1^2^*-40, *drp1^1^*-59, *drp1^KG^*- 67) **E)** Top: Immunoblot analysis of anterior region of *Drosophila* ovarioles (germarium and stages 2-3) following germline-specific knockdown of Drp1, or Opa1; numbers signify levels normalized (n.) with mCherry RNAi for each lane. Bottom: Representative confocal optical slice of AtpB immunostained GSCs expressing mCherry, Drp1, or Opa1 RNAi using germline-specific Nanos- GAL4 driver; scale bar : 2 μm **F)** Boxplots showing egg-production on Day 2 and 12 following germline-specific knockdown of mCherry, Drp1, or Opa1. (Day2-N-Basal: mCherry RNAi-3, Drp1 RNAi-3, Opa1 RNAi-3; Day12- N-Basal: mCherry RNAi-28, Drp1 RNAi-17, Opa1 RNAi-17; Day2-N-Stimulated: mCherry RNAi- 3, Drp1 RNAi-3, Opa1 RNAi-3; Day12-N-Stimulated: mCherry RNAi-22, Drp1 RNAi-27, Opa1 RNAi-13) **G)** Boxplot showing egg-production on Day 12 following germline-specific knockdown of mCherry, Drp1, or Opa1 and simultaneous knockdown of Drp1 and Opa1 (N-Basal: mCherry RNAi-10, Drp1 RNAi-14, Opa1 RNAi-15, Drp1 and Opa1 RNAi-15 ; N-Stimulated: mCherry RNAi-10, Drp1 RNAi-13, Opa1 RNAi-14, Drp1 and Opa1 RNAi-13 ) **H)** Representative confocal micrographs of germaria immunostained for pH3 and Hoechst DNA stain following germline-specific knockdown of mCherry, Drp1, or Opa1; scale bar : 5 μm **I)** Bar plots showing abundance of pH3-positive cells (%) in Region 1 of stimulated (12 days) germaria following germline-specific knockdown of mCherry, Drp1, Opa1, and simultaneous Drp1 and Opa1 knockdown. (N : 15 germaria for each genotype) **J)** Boxplots showing egg-production on Day 12 following germline-specific knockdown of mCherry, Drp1 and Opa1 in CD and HSD (N : 15 for each genotype) **K)** Bar plots showing abundance of pH3-positive cells in Region 1 of germaria following germline- specific knockdown of mCherry, Drp1 and Opa1 in CD and HSD (N : 15 germaria for each genotype) **L)** Line plots showing GSC Self-renewal / Proliferation (black) and Loss (grey-dashed) in CD and HSD. **M)** Bar plots showing abundance of FSCs (%) with >1 division per cyst in CD and HSD. (N(cyst)- CD: WT-81, *drp1^2^*-33, *drp1^1^*-59, *drp1^KG^*-27; N(cyst)-HSD: WT-41, *drp1^2^*-52, *drp1^1^*-42, *drp1^KG^*-24)

The FSC lineage tracing analyses showed that the abundance of clonal *drp1^2^ and drp1^1^*FSCs is low due to their increased loss (Fig. S1F). Notably, the *drp1^KG^* FSCs exhibit prominently elevated self-renewal / proliferation with the dietary stimulation (Fig. 1D, inset showing abundance of FSCs with >1 divisions). Therefore, reduction of Drp1 boosts FSC self-renewal / proliferation towards differentiating them into FCs, while the GSC and FSC functionality depends on different levels of Drp1. This difference on Drp1 dependency could be due to underlying differences in cell cycle, given Drp1 functionality is integrated in cell cycle regulation (Taslim et al. 2023; Spurlock et al. 2020). Indeed, the cell cycle Fly-FUCCI reporter indicates that the FSCs are not in the G2- M(pre-mitosis) state where the majority of the GSCs reside (Fig. S1G); the proliferating germline cells go through S (DNA synthesis) and reside at G1(long growth) after differentiation.

To assess the impact of Drp1-tuning driven boost in ovarian stem cell functionality on egg production, we expressed RNAis against Drp1 or Opa1 in the GSC (using Nanos-Gal4) and FSC lineages (using 109-30-Gal4) to attenuate their mitochondrial fission or fusion activity, respectively; mCherry-RNAi is used as control. Levels of successful knockdown for both Drp1 and Opa1 confirm that Drp1 repression is incomplete to allow its fine-tuning, which did not show a visible change in mitochondrial size while the Opa1-RNAi visibly reduced mitochondrial size (Fig. 1E). Egg production due to stimulation of ovarian stem cells by protein-rich dietary supplement is detected after 10-12 days of egg development, while that due to direct nutritional influences is detected within 2 days (Bastock and St Johnston 2008; Hsu and Drummond-Barbosa 2017). Notably, GSCs-lineage specific incomplete Drp1-knockdown boosts and that with Opa1- knockdown suppress egg production with the dietary stimulation (12 days), compared to the control (Fig. 1F); other Drp1/Opa1 RNAis exhibit similar impacts (Fig. S1H). Such impacts are also detected in basal conditions over 2 and 12 days, demonstrating protein supplementation elevates the baseline impact of the knockdowns. Importantly, the opposite impact of the GSCs-lineage specific Drp1 and Opa1 knockdown on egg production is countered in both basal and stimulated conditions by combinatorial expression of Drp1 and Opa1 RNA-i, which does not happen with mCherry-RNAi (Fig 1G, Fig. S1I). Consistently, stimulated Drp1-knockdown GSCs and their progenies in the Region 1 (upto 4 cell-stage) exhibit pronounced increase in pH3 positive cells (mitotic marker), in comparison to the control (Fig 1H). These data suggest that Drp1-tuned GSCs and their progenies have higher mitotic rate and/or length in dietary stimulated condition, consistent with enhanced self-renewal detected in the *drp1^2^*-GSCs (Fig. 1B). Moreover, Opa1 knockdown decreases the abundance of the pH3 positive cells and the combinatorial knockdown with Drp1-RNAi normalizes it (Fig. 1I, Fig. S1I). Therefore, these data suggest that tuned Drp1 repression-driven boost of GSC self-renewal/proliferation leading to enhanced egg production is brought about by mitochondrial regulation involving Opa1. Drp1 tuning by its partial repression (in a hypomorphic mutant or by RNAi) allows Drp1 activation. Indeed, GSC-lineage specific Drp1 overexpression also increases pH3 positive cells in region 1 in comparison to RFP control, without boosting egg production likely due to defects during egg development (Fig. S1J). FSC-lineage specific Drp1 knockdown does not show any boost of egg production in basal or stimulated condition, consistent with prominent positive impact only with the strongest Drp1 mutant (Fig. S1K). Nonetheless, knockdown of Opa1 in the FSCs suppresses egg production, indicating the Opa1 dependency is similar between GSCs and FSCs (Fig. S1K).

Thus, data from gene-diet interaction studies employing two genetic strategies demonstrate that *Drosophila* ovarian stem cells are tuned by Drp1 driven mitochondrial regulation involving Opa1 to prime their response to protein-rich supplementation. The optimal Drp1 level/activity of the primed stem cell state allows protein-rich supplementation to enhance their self-renewal, proliferation and differentiation to boost egg production. Also, the data highlights that different levels of Drp1 activity support optimal stem cell activity in different stem cell lineages even within the same tissue.

### Drp1 driven priming of *Drosophila* ovarian stem cells reverses the inhibitory impact of high sugar diet (diabetes-like condition) on egg development, with protein-rich dietary supplement

High sugar diet formulation for *Drosophila* induces the systemic hyperinsulinemic-hyperglycemic environment of diabetes and profoundly inhibits egg production (Nunes and Drummond-Barbosa 2023; Palanker Musselman et al. 2011). *Drosophila* on High Sugar Diet (HSD) for 8 days have elevated fat mass in comparison to control diet (CD), thus reflecting the systemic metabolic changes (data not shown). Here we investigated if and how Drp1 regulated mitochondrial priming of GSCs/FSCs modulates the HSD impact on egg development and production over 12 days.

The impact of additional protein-rich supplementation is less in CD than in a corn meal diet due to the double basal protein content of CD (see Methods). Similarly, the stimulatory impact of GSC-lineage specific Drp1-knockdown on egg production is only modest above the control (Fig. 1J). Nonetheless, HSD reduces basal egg production from control and incomplete Drp1- knockdown GSCs. Whereas with protein-rich stimulation, GSCs-lineage specific incomplete Drp1-knockdown (primed) completely rescues the egg production defect of HSD. This fails to happen with the GSCs-lineage specific Opa1-knockdown (defective priming), despite HSD boosting basal egg production from them modestly but significantly. These data suggest that the mitochondria primed GSCs and/or their progenies depend on protein-rich supplement, and on HSD environment when priming is defective. Surprisingly, HSD elevates mitosis (rate and/or length) of the control GSCs and their progenies similar to protein-rich supplement, as revealed by the abundance of pH3 positive nuclei in region 1 (Fig. 1K). This suggests HSD inhibition of egg production is likely due to its negative impact on the later stages of egg development. Also, HSD maintains the stimulated mitosis in Drp1-knockdown GSCs (primed) and the inhibited mitosis of Opa1-knockdown GSCs (defective in priming), similar to CD. Protein-rich supplementation in HSD does not further stimulate the control or knockdown GSCs. Notably, combination of HSD and protein-rich supplementation is most inhibitory for Opa1-knockdown GSC mitosis and egg production.

Next, we investigated the impact of Drp1 tuning on GSC self-renewal and differentiation in the presence of HSD and protein-rich supplementation, employing the lineage tracing analyses of the control and Drp1 mutant GSCs / FSC clones (as in Fig. 1A-B). Abundance of clones varies strikingly between CD and HSD primarily due to the loss of clones in the HSD group (Fig. 1L, Fig. S1L-M). More importantly, consistent with elevated mitosis of the control GSCs (Fig. 1K), HSD increases self-renewal/proliferation and differentiation of the WT-GSCs (Fig 1L, Fig. S1L-N). However, *drp1^2^*-GSCs balance the loss and self-renewal under the influence of the HSD (Fig. 1L, Fig. S1M), while their progeny cysts appear larger than other groups (Fig. S1O). Therefore, the HSD driven inhibition of egg production in control GSCs appears to be primarily due to loss of the germline cysts that is reversed by Drp1 fine-tuning, thus explaining the reversal of the HSD defect. Nonetheless, HSD does not alter the self-renewal of the WT-FSCs, but increases their loss (Fig. 1M, Fig. S1P-Q). More importantly, the maximum proliferation of the *drp1^KG^* -FSCs stimulated in CD is further markedly enhanced by HSD (Fig. 1M, S1Q).

In summary, HSD does not inhibit self-renewal, proliferation and differentiation of the normal or primed *Drosophila* ovarian stem cells but reduces egg production due to increased loss of the egg chambers. Such a growth defect is overcome by the priming of GSCs by tuned repression of Drp1. The data highlights the difference in metabolic dependencies between GSC and FSCs and indicates that HSD may provide a proliferative environment for the somatic FSCs when Drp1 is almost completely repressed.

### Precise tuning of Drp1 allows its characteristic recruitment and organization on mitochondria in primed *Drosophila* ovarian GSCs with mildly reduced Drp1 activity

To elucidate the mechanism of mitochondrial priming of *Drosophila* ovarian GSCs, we probed for any unique quantitative mitochondrial characteristics of tuned repression of Drp1 in the primed *drp1^2^*-GSCs, given complete loss of Drp1 is inhibitory for stemness and other cellular properties (Favaro et al. 2019; Ishihara et al. 2009; Spurlock et al. 2021b). Immunohistochemistry expectedly detects markedly reduced Drp1 signal in the homozygous *drp1^KG^* functional null clones in comparison to the heterozygous WT background (Fig S2A). Therefore, we used this approach to compare the functional levels and quantitative characteristics of the Drp1 protein in the WT and Drp1 mutants. Indeed, the ratio of immuno-stained Drp1 signal in the clonal GSCs to the adjacent non-clonal GSCs is pronouncedly lower in the *drp1^KG^*-GSCs in comparison to the *drp1^1^, drp1^2^* or WT-GSCs (Fig. S2B).

Next, we developed and validated an image analysis pipeline to identify the quantitative characteristics of Drp1 ‘on’ or ‘not on’ the resolvable mitochondrial units in each GSC by near-super resolution AiryScan2 microscopy. To characterize and quantify mitochondrial-units and Drp1 clusters in clonal GSCs, a connected-component analysis pipeline was developed using CellProfiler and MATLAB on original deconvolved GSC Z-stacks. Mitochondria-associated Drp1 pixels were defined by exact spatial overlap with mitochondrial pixels. Subsequently, both the mitochondrial and Drp1 channels were evaluated via connected-component analysis to compute multiple features (Fig 2A, S2C). Validation of this analysis comes from the progressive decrease in total number of Drp1 pixels and the fraction overlapping with mitochondrial pixels in individual clonal GSCs with decreasing Drp1-activity (Fig S2D, E). Consistently, there is progressive increase in the fraction of mitochondrial-units with ‘0’ Drp1-clusters with decreasing Drp1 activity (Fig. 2B, inset). Moreover, bivariate analysis of the ‘size’ of individual mitochondrial-units and the total ‘number’ of their Drp1-clusters (0 and above) confirmed that the mitochondrial-units of the strongest *drp1^KG^* mutant have the least number of Drp1 clusters irrespective of mitochondrial-unit size (Fig. 2B). Interestingly, the number of mitochondrial Drp1-clusters increases exponentially with mitochondrial-unit size beyond a certain threshold (Fig. 2B).

**Figure 2.**
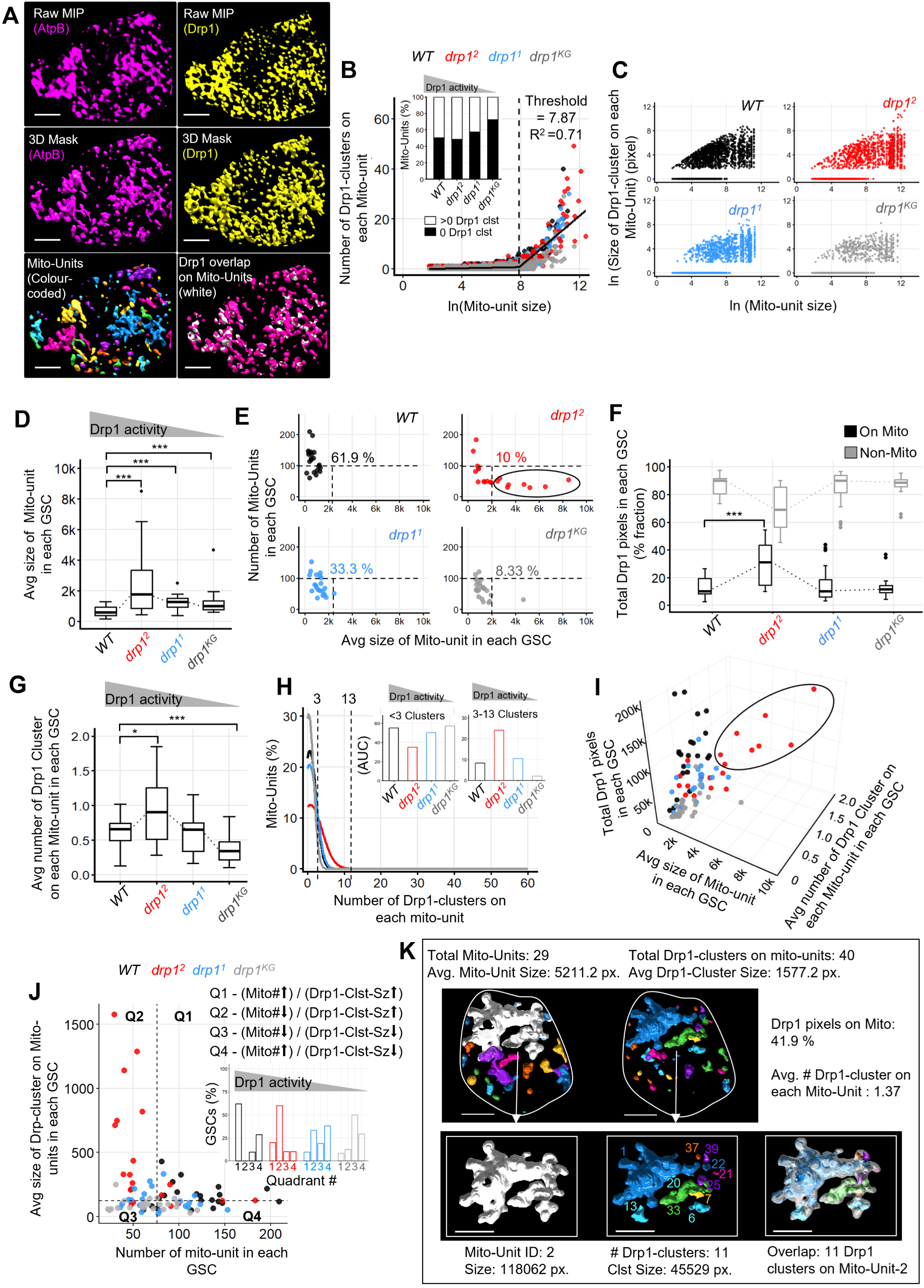
**A**) Representative MIP and a section of 3D rendered micrograph of WT-GSC clone showing mitochondrial-units and Drp1-clusters generated by connected-component analysis (See Methods). Raw maximum intensity projections (MIP) get converted to 3D Masks to obtain the color coded mitochondrial-units and the Drp1 overlapping pixels on individual mitochondrial-units. **B)** Bivariate scatter plot of mitochondrial-units of stimulated GSCs for each genotype showing relationship between their log normalized (ln) size and total number of Drp1-clusters on them. Inset showing stacked-barplots of mitochondrial-units (%) with or without Drp1 clusters; gray triangle depicting decreasing Drp1 activity across the mutants. (N(mitochondrial-units) : WT-2477, *drp1^2^*-1313, *drp1^1^*-1655, *drp1^KG^*-1582). **C)** Bivariate scatter plot of mitochondrial-units of stimulated GSCs for each genotype showing relationship between their log normalized (ln) size and (ln) size of individual Drp1-clusters on them (N(Drp1-clusters) : WT-3085, *drp1^2^*-1965 , *drp1^1^*-2146 , *drp1^KG^*-1819). **D)** Boxplots showing average size of mitochondrial-units in each stimulated GSC for each genotype (N(GSC) : WT-21, *drp1^2^*-20, *drp1^1^*-22, *drp1^KG^*-24). **E)** Bivariate scatter plot of stimulated GSCs for each genotype showing the relationship between the total number and average size of their mitochondrial-units (N(GSC) : WT-21, *drp1^2^*-20, *drp1^1^*- 22, *drp1^KG^*-24). **F)** Boxplots showing fraction of Drp1-pixels (%) recruited (black) or not-recruited (gray) on mitochondria in each stimulated GSC of each genotype (N(GSC) : WT-21, *drp1^2^*-20, *drp1^1^*-22, *drp1^KG^*-24); dotted lines connect the medians. **G)** Boxplots showing average number of Drp1-clusters on each mitochondrial-unit in each stimulated GSC of each genotype (N(GSC) : WT-21, *drp1^2^*-20, *drp1^1^*-22, *drp1^KG^*-24) **H)** Frequency distribution (%) of mitochondrial-units with a given number of Drp1-clusters in each genotype with dashed lines representing 3 and 13 Drp1-clusters on each mitochondrial-unit. Inset bar plot showing area under the curve (AUC) corresponding to mitochondrial-units with <3 and 3–13 characteristic Drp1-clusters. (N(mitochondrial-units) : WT-2477, *drp1^2^*-1313 , *drp1^1^*-1655, *drp1^KG^*-1582) **I)** Trivariate scatter plot of stimulated GSCs showing the relationship among the average size of their mitochondrial-units, average number of Drp1-clusters on each mitochondrial-unit, and total number of Drp1 pixels detected in each GSCs. (N(GSC) : WT-21, *drp1^2^*-20, *drp1^1^*-22, *drp1^KG^*-24) **J)** Bivariate scatter plot of stimulated GSCs of each genotype showing relationship between their total number of mitochondrial-units and the average size of their Drp1-clusters, where quadrants are demarcated with Median Absolute Deviation (MAD) values. Inset bar plot showing the % GSCs from each genotype distributed across the quadrants. (N(GSC) : WT-21, *drp1^2^*-20, *drp1^1^*- 22, *drp1^KG^*-24). **K)** Representative section of 3D rendered micrograph of mitochondrial-units and mitochondrially- recruited Drp1-clusters of a drp1^2^-primed-GSC from the highlighted GSC subpopulation in 2E / I. The mitochondrial-unit with the largest size and its associated overlapping Drp1-clusters are highlighted, with the corresponding values of their characteristic properties mentioned.

Mitochondrial-unit size is expected to increase with unopposed mitochondrial fusion. Interestingly, the maximum ‘size’ of the mitochondrial Drp1-clusters increases with the average ‘size’ of the mitochondrial-unit that is maximum in the *drp1^2^-*GSCs and modest but significantly higher in the other two Drp1 mutants (Fig. 2C-D). Next, we probed into bivariate analyses of the average ‘size’ and ‘number’ of the mitochondrial-units in individual GSCs. Expectedly, more WT- GSCs are detected with >100 mitochondrial-units than the *drp1* mutant GSCs that allow unopposed fusion of individual mitochondrial-units. Uniquely, a *drp1^2^-*GSC subpopulation harbors <100 mitochondrial-units that are >2000 pixels in size (Fig. 2E, outlined). This data highlights the plasticity of the structure of the mitochondrial-units between *drp1^2^-*GSC subpopulations, which may arise from tunability of Drp1 protein with mildly reduced function.

Therefore, we investigated if the uniquely large mitochondrial-units of the primed *drp1^2^-* GSCs have any characteristic Drp1 recruitment and organization on them. Surprisingly, *drp1^2^-* GSCs have an elevated fraction of mitochondrial-Drp1 clusters and lower non-mitochondrial fraction, in comparison to other GSCs (Fig. 2F). Indeed, the average number of Drp1 clusters per mitochondrial-unit is highest in the *drp1^2^-*GSCs (Fig. 2G). Careful analysis of the frequency distribution of the mitochondrial-units with a given number of Drp1 clusters revealed that the *drp1^2^-* GSCs have maximum abundance of mitochondrial components with 3 to 13 detectable Drp1 clusters and minimum abundance of those with less than 3 Drp1 clusters (Fig. 2H and inset). A trivariate analysis of the above mitochondrial and Drp1 characteristic features confirmed that the greater number of Drp1-clusters are on the unique larger mitochondrial-units of the *drp1^2^-*GSC subpopulation with intermediary Drp1 levels (Fig. 2I, outline). Given mitochondrial fission is expected to increase the number of mitochondrial components, we finally performed bivariate analyses of the ‘total-number’ of mitochondrial-units and ‘mean-size’ of mitochondrial Drp1- clusters in individual GSCs across the genetic groups. MAD based analysis (see Methods) of the bivariate scatter plot identified 4 GSC groups. Most uniquely, >60% of the *drp1^2^-*GSCs have fewer mitochondrial-units with no WT-GSCs demonstrating these characteristics (Fig. 2J and inset).

Therefore, our quantitative analyses revealed that mild reduction of Drp1 activity, as in *drp1^2^-*GSCs, allows a characteristic pattern of Drp1 recruitment and organization on mitochondria, linked to their enhanced priming properties. Particularly, a unique subpopulation of *drp1^2^-*GSCs exhibits enhanced Drp1 recruitment on larger mitochondrial-units that are fewer in number (represented in Fig. 2K). This may result from unopposed Opa1 driven mitochondrial fusion allowed by tuned lowering of Drp1 activity, given they genetically counter each other. It is important to note that the absolute number and size of Drp1 is limited at the level of near-super resolution AiryScan2 microscopy, which may mask minor differences within the comparative scale.

### Characteristic Drp1 recruitment and organization on mitochondria is dependent on interaction of Cry driven rhythm and protein-rich supplementation in *Drosophila* ovarian GSCs

The priming response to protein-rich supplementation for increasing egg production is enhanced by mild genetic repression of Drp1 in the GSCs that maintains a characteristic Drp1 recruitment and organization on mitochondria, as studied within a 24-hour light-dark cycle (Fig. 1-2). Here, we investigate if the control GSCs exhibit similar dietary regulation of Drp1 under the influence of light and dark components. Therefore, we studied GSC priming properties of Drp1 in mid-light / mid-dark, with and without protein-rich supplement, in control and mutants with perturbed rhythm. We used W^1118^ control, and mutants of the Cry protein, Cry^b^ (deficient in light sensing) and Cry^out^ (gene ablated) that link the diurnal and the circadian rhythms (Stanewsky et al. 1998; Damulewicz and Mazzotta 2020; Yoshii et al. 2008).

Basal Drp1 protein levels undergo oscillations between mid-light and mid-dark in the early-stage *Drosophila* egg chambers (germarium and stages 2-3), consistent with findings in other tissues in mice (Collins et al. 2021; Schmitt et al. 2018). Specifically, Drp1 level is prominently lowered in mid-dark than in mid-light in W^1118^, when ATP-B level (mitochondrial marker) does not follow the same rhythmicity (Fig. 3A). This rhythmicity of Drp1 protein is lost in both Cry mutants, as evident in their constitutively high Drp1 levels (Fig. 3A). The basal Drp1 level per unit mitochondrial content in GSCs, measured as ratio of Drp1/AtpB immunostain signal decreases as W^1118^ or Cry^out^ GSCs give rise to the proliferating 4-cell-cystoblasts that terminally differentiate into the 16-cell-cysts (Fig. 3B-C, S3A). This decrease in Drp1 levels along differentiation is not observed in the FSC lineage, at least within the germarium. Nonetheless, the oscillation of Drp1 per unit mitochondrial content between mid-light-high and mid-dark-low levels is perturbed in the Cry^out^-GSCs/ FSCs and their progenies (Fig. 3B-C, S3A). Thus, this data corroborates the observation from the biochemical analyses of the whole tissue (Fig. 3A). Stimulation with protein-rich supplement (12 days) prominently elevates the baseline Drp1 levels per unit mitochondrial content in the W^1118^-GSCs, while maintaining the oscillating mid-light-high and mid-dark-low levels (Fig. 3D, S3B); this is detectable within 2 days, as shown in the WT Canton S GSCs (Fig. S3C). Notably, the baseline elevation of Drp1 levels is dampened in the Cry^out^-GSCs that maintains the defect in Drp1 rhythmicity (Fig. 3D, S3B).

**Figure 3.**
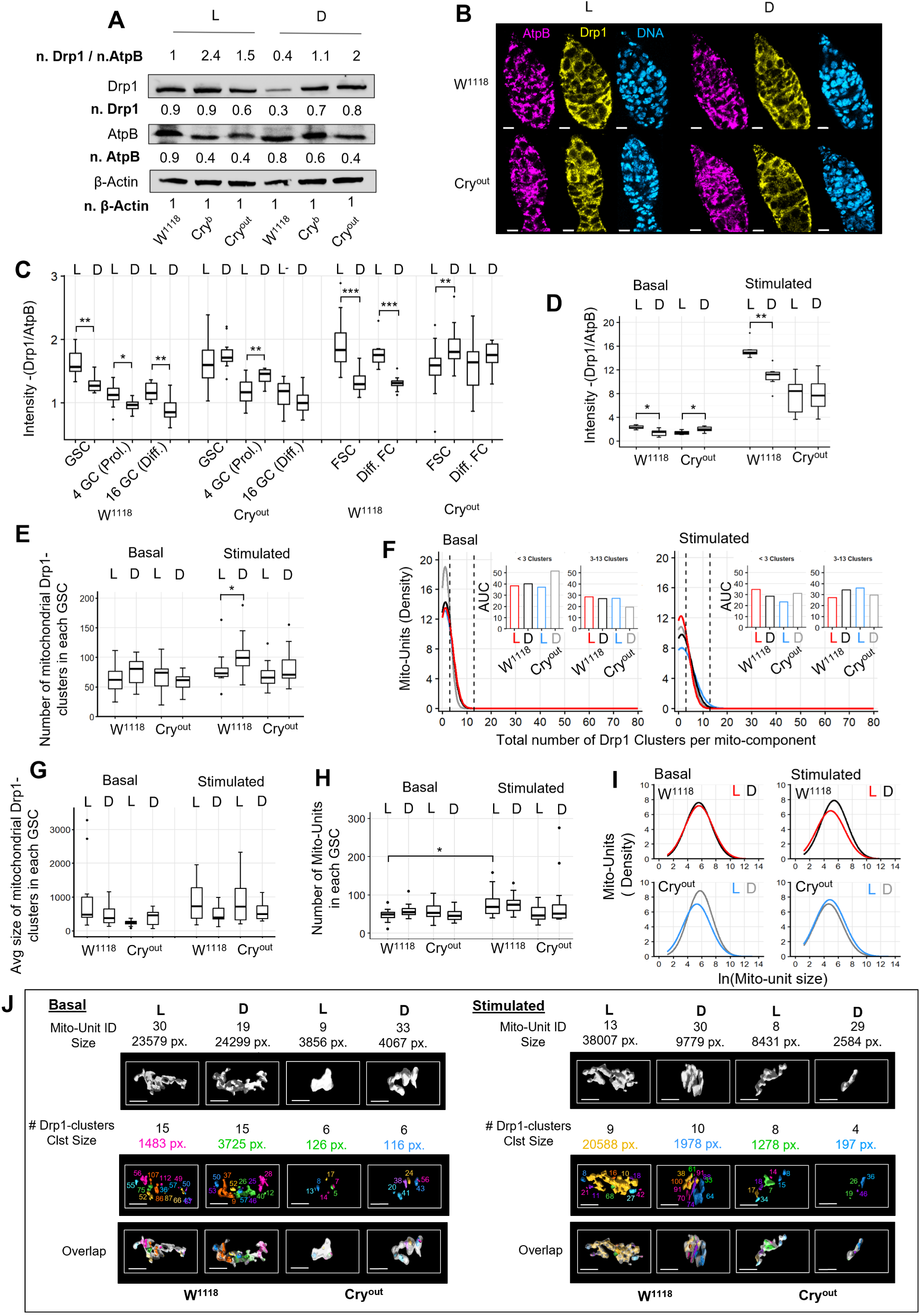
**A**) Immunoblot analysis for the Drp1 and AtpB in W^1118^ ,Cry^b^ and Cry^out^ mutant ovaries in mid- light(L) and mid-dark(D) in basal condition; numbers signify levels normalized (n.) with β-Actin for each lane. B) Representative confocal optical slice of W^1118^ and Cry^out^ germaria in mid-light(L) and mid- dark(D) in basal condition immunostained for AtpB and Drp1, with Hoechst DNA stain. C) Boxplots showing the Drp1 intensity normalized by AtpB in GSC and FSC lineages in the germaria of W^1118^ and Cry^out^ in mid-light(L) and mid-dark(D) in specified developmental stages in basal condition, as quantified from experiment described in B. (N-Basal-10/cell type/genotype/time point; except FSCs_N-20) D) Boxplots showing the Drp1 intensity normalized by AtpB intensity in GSCs of W^1118^ and Cry^out^ germaria in mid-light(L) and mid-dark(D) in basal and stimulated conditions. (N- 6/genotype/timepoint/diet) E) Boxplots showing number of Mitochondrial Drp1-clusters in each GSC (N(GSC)- 12/genotype/timepoint/diet) F) Frequency distribution (%) of mitochondrial-units with a given number of Drp1-clusters in each genotype with dashed lines representing 3 and 13 Drp1-clusters on each mitochondrial-unit. Inset bar plot showing area under the curve (AUC) corresponding to mitochondrial-units with <3 and 3–13 characteristic Drp1-clusters. (N(mitochondrial-units)-Basal : W^1118^- L-610 ,D-709; Cry^out^- L- 721, D-587; N(mitochondrial-units)-Stimulated : W^1118^ - L-939, D-924; Cry^out^ - L-627 , D-988) G) Boxplot of Average size of mitochondrial Drp1-clusters in each GSC (N- 12/genotype/timepoint/diet) H) Boxplot of # Mito-Units in each GSC (N-12/genotype/timepoint/diet) I) Frequency distribution (density) of mitochondrial-units with their log normalized (ln) size. (N(mitochondrial-units)-Basal: W^1118^- L-610 ,D-709; Cry^out^- L-721, D-587; N(mitochondrial-units)- Stimulated: W^1118^ - L-939, D-924; Cry^out^ - L-627 , D-988) J) Representative section of 3D rendered micrograph of characteristic mitochondrial-unit and its associated overlapping Drp1-clusters of W^1118^- and Cry^out^-GSCs in mid-light(L) and mid-dark(D) The corresponding values of their characteristic properties mentioned.

Next, we studied the priming characteristics of Drp1 protein (as in Fig. 2), focusing on the W^1118^/ Cry^out^ GSCs. The basal Cry^out^ GSCs harbor more mitochondrial-units without Drp1-clusters particularly in mid-light although the total detected Drp1 pixels is higher, indicating a defect in Drp1 recruitment in the absence of Cry (Fig. S3D-E). Moreover, stimulated Cry^out^-GSCs fail to increase the number of mitochondrially recruited Drp1 clusters in mid-dark observed in the W^1118^- GSCs despite their lower Drp1 levels (Fig. 3E). Probing into the mitochondrial Drp1-clusters revealed that the mitochondrial population with characteristic 3 to 13 Drp1-clusters per mitochondrial-unit of the primed *drp1^2^-*GSCs is modestly increased only in mid-dark in the stimulated W^1118^-GSCs (Fig. 3F). Contrarily, the basal defect of Cry^out^-GSCs is reflected in the higher abundance of mitochondrial population with <3 Drp1 clusters, while the stimulated Cry^out^- GSCs exhibit an overall increase in Drp1 recruitment in both mid-light and mid-dark (Fig. 3F). Moreover, the size of the mitochondrial Drp1-clusters of the basal Cry^out^-GSCs in mid-light is dramatically small compared to that of W^1118^-GSCs, and stimulation increases it in both W^1118^/ Cry^out^ GSCs (Fig. 3G) that is likely due to elevated Drp1 levels (Fig. 3D).

The larger Drp1-clusters may represent the mature fission-active Drp1 puncta that arise from the immature smaller Drp1 puncta in cell culture models (Ji et al. 2015; Rosenbloom et al. 2014). Active mitochondrial fission in the stimulated W^1118^-GSCs of mid-light-cycle is reflected in their greater average Drp1-cluster size, higher mitochondrial number and reduced mitochondrial- unit size in comparison to those of mid-dark-cycle (Fig. 3G-I). These characteristic features are also observed in the stimulated wild type GSC clones, while the fission defect in the *drp1^2^-*GSCs sustains the large Drp1-clusters on fewer large mitochondrial-units (Fig. 2E, J). Therefore, our consistent data suggest that large-mature Drp1-clusters drive mitochondrial fission to maintain greater number of small mitochondria in mid-light and the small-immature Drp1-clusters allow unopposed mitochondrial fusion in mid-dark to maintain fewer larger mitochondria in the stimulated W^1118^-GSCs. On the other hand, in the basal Cry^out^-GSCs, the mitochondrial-unit size remains smaller in mid-light (Fig. 3I), where a greater number of small Drp1 clusters are recruited (Fig. 3F-G). This defect may disallow the dietary stimulation driven regulation of Drp1 and mitochondria observed in W^1118^-GSCs (Fig. E-I). Thus, Cry protein is directly or indirectly involved in regulation of mitochondrial Drp1 properties under protein-rich supplementation. The canonical Cry function is exerted in the brain, and Cry has not been detected in *Drosophila* ovary with classical techniques (Rush et al. 2006; Emery et al. 2000). However, Cry transcripts can be clearly detected in Drosophila ovarian cells with more sensitive detection methods like single cell transcriptomics from two independent studies (Fig. S3F) (Jevitt et al. 2020; Rust et al. 2020). Consistently, Cry-Gal4 is able to drive RFP in both GSC and FSC lineages demonstrating active Cry promoter in these lineages (Fig. S3G). It remains to be tested if the prominent impact of Cry on Drp1 in the *Drosophila* ovarian stem cells can be possibly exerted directly in these cells.

Thus, Cry dependent light-dark components interact with protein-rich supplementation to modulate Drp1 recruitment and organization in normal GSCs (represented in Fig. 3J, Fig. S4A). In the dark cycle, the reduced level of Drp1 is stimulated by the protein-rich supplement to be recruited onto mitochondria in characteristic numbers and reduced cluster size of restricted fission ability. In the light-cycle with higher Drp1 protein level, the Drp1-clusters mature in size when stimulated by protein-rich supplement to trigger mitochondrial fission. Such characteristic recruitment and organization of Drp1 with associated alteration in mitochondrial unit-size is perturbed in the absence of Cry that maintains constitutively high levels of Drp1. The reduced Drp1 levels of the dark cycle recapitulates reduced Drp1 activity of the primed *drp1^2^-*GSCs, raising the possibility that mitochondrial priming of normal GSCs is initiated in the dark-cycle.

### Drp1 driven mitochondrial priming of Drosophila ovarian GSCs and egg production is dependent on interaction between Cry function and dietary Threonine levels

We investigated if the described interaction of light-dark rhythm and dietary components for maintaining characteristic priming properties of the mitochondrial Drp1 pool (Fig. 3) lead to GSC priming driven enhanced egg production. Protein-rich dietary supplement accelerates entrainment of egg production to light-dark (LD, normal), constant light (LL, perturbed by constant Cry activation) or constant dark (DD, free running) (Dubowy and Sehgal 2017) regimens (not shown). In entrained WT Canton S strain, egg production has the following attributes (Fig. S4B): **a)** higher in light-cycle than in dark-cycle in basal and dietary stimulated conditions; **b)** stimulation from basal (over 12 days) is detected most prominently in dark-cycle, and only in light-cycle in DD free-running rhythm; **c)** light-dark difference is dampened in LL with constant Cry activation.

Next, we studied the influence of LL and DD regimens on Drp1-driven regulation of GSC properties in basal and dietary stimulated conditions. We performed lineage tracing analysis using the *drp1* mutants, followed by analyses of egg production and pH3 positive GSCs and their progenies after GSC-specific knockdown of Drp1 (as in Fig.1 for LD). Interestingly, no germline clones of the *drp1^KG^* functional null mutants are detected in LL or DD (Fig. S4C), consistent with their reduced clonal abundance in LD (Fig. S1D); clonal abundance of the other hypomorphic mutants are comparable to LD. This indicates Drp1 functionality being essential for GSC adaptation to environmental light-dark cycles. While stimulation of self-renewal/proliferation of the W^1118^-and *drp1^1^* GSCs is more prominent in LL than in DD, such conditions suppress the enhanced mitochondrial priming of the *drp1^2^*-GSCs detected previously in LD (Fig. 4A-B, Fig. 1A- B). Consistently, the Drp1 knockdown driven boost in egg production is also not detected when stimulated in LL and DD (Fig. 4C). However, the Drp1-knockdown GSCs and their progenies still sustain mitosis (rate or length) at a subjective light-cycle when stimulated in LL and DD, maximally in the latter (Fig. 4C, inset). This likely reflects proliferation of differentiated cystoblasts and not self-renewal, as primed *drp1^2^*-GSCs sustain differentiation only when stimulated in DD (Fig. S4D). On the other hand, the W^1118^-GSCs and their progenies exhibit marked suppression of mitosis (rate or length) when stimulated in DD (Fig. 4C, inset), which may contribute to their reduced egg production in the dark-cycle (Fig. S4A). Therefore, mitochondrial priming of GSCs requires cycling through light and dark.

**Figure 4.**
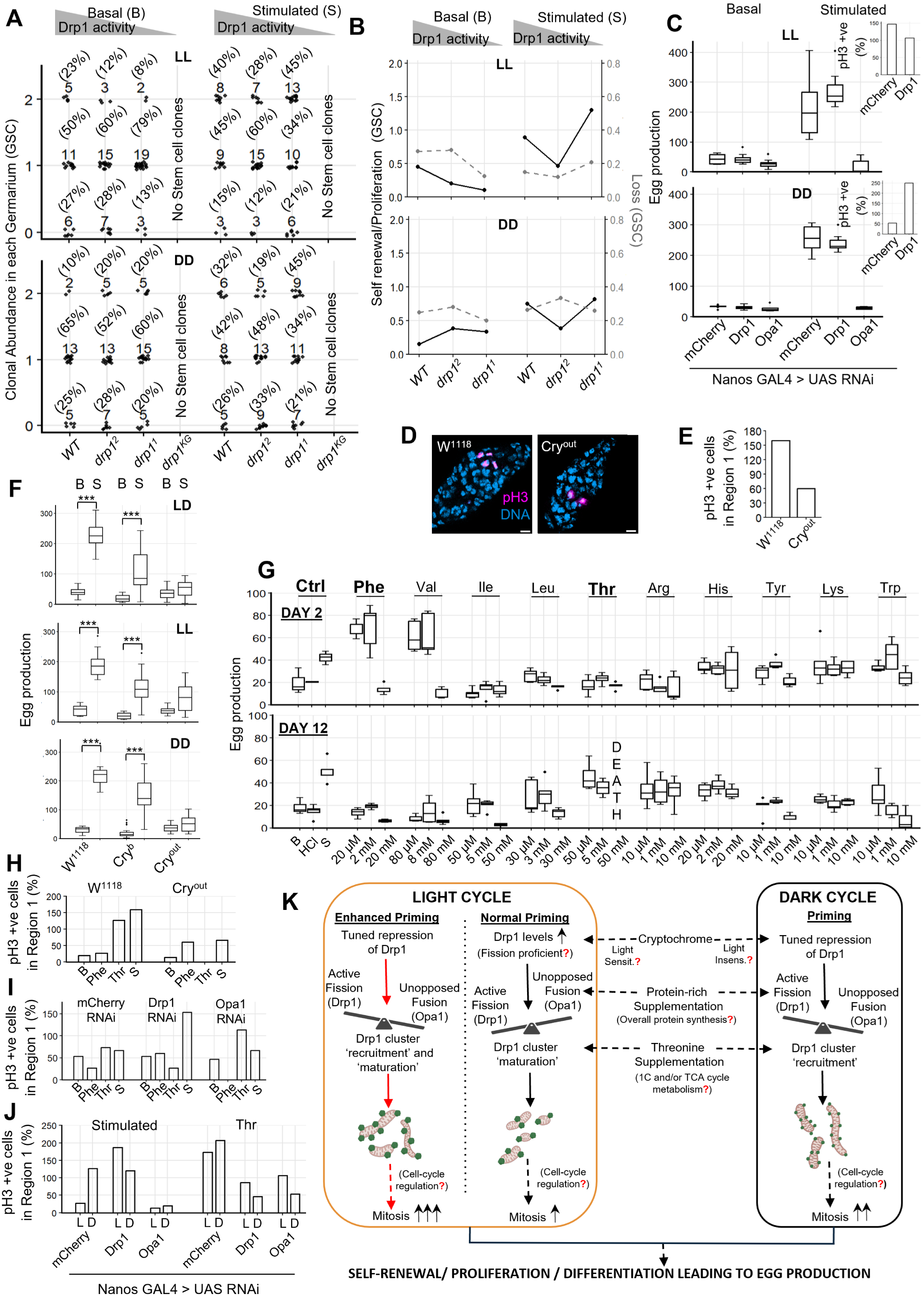
**B**) Dot plots showing clonal GSC abundance in each germarium with 0-, 1-, and 2-GSCs where frequency (%) denoted in parentheses for each genotype in LL and DD conditions; gray triangle depicting decreasing Drp1 activity across the mutants. Stem cell clones were undetected in *drp1^KG^*(N(germaria)-Basal: LL - *WT*-22, *drp1^2^*-25, *drp1^1^*-24; DD - WT-20, *drp1^2^*-25, *drp1^1^*-25; N(germaria)-Stimulated : LL - *WT*-20, *drp1^2^*-25, *drp1^1^*-29; DD - *WT*-19, *drp1^2^*-27, *drp1^1^*-27) C) Line plots showing GSC Self-renewal / Proliferation (black) and Loss (grey-dashed), as calculated from A in LL and DD D) Boxplots showing egg-production on Day 12 following germline-specific knockdown of mCherry, Opa1 and simultaneous knockdown of mCherry and Opa1 in LL and DD. (N-Basal: LL - mCherry RNAi-10, Drp1 RNAi-10, Opa1 RNAi-10; DD- mCherry RNAi-9, Drp1 RNAi-10, Opa1 RNAi-5; N-Stimulated: LL - mCherry RNAi-10, Drp1 RNAi-9, Opa1 RNAi-10; DD- mCherry RNAi- 10, Drp1 RNAi-9, Opa1 RNAi-5). Inset shows abundance of pH3-positive cells (%) in Region 1 of germaria (N : 15 germaria for each genotype). E) Representative confocal micrographs of W^1118^ and Cry^out^ germaria immunostained for pH3 and Hoechst DNA stain; scale bar : 5 μm. F) Bar plots showing abundance of pH3-positive cells (%) in Region 1 of W^1118^ and Cry^out^ germaria (N : 30 germaria for each genotype) G) Boxplots showing egg-production on Day 12 in W^1118^ ,Cry^b^ and Cry^out^ in basal and stimulated conditions in LD, LL and DD regimes. (N : W^1118^ - 15/diet; Cry^b^ and Cry^out^ -20/diet) H) Boxplot showing egg-production on Day 2 and Day 12 in W^1118^ supplemented with essential amino acids (N: Controls- 3/diet; Amino acids- 5/Amino acid conc.) I) Bar plots showing abundance of pH3-positive cells (%) in Region 1 of W^1118^ and Cry^out^ germaria (N: 15 germaria for each diet for each genotype) J) Bar plots showing abundance of pH3-positive cells (%) in Region 1 of germaria following germline-specific knockdown of mCherry, Drp1, Opa1 (N: 15 germaria for each diet for each genotype) K) Bar plots showing abundance of pH3-positive cells (%) in Region 1 of germaria following germline-specific knockdown of mCherry, Drp1, or Opa1 in yeast- and threonine-supplemented groups in mid-light(L) and mid-dark(D) (N : 15 germaria for each genotype). L) Schematic model depicting Drp1-dependent priming under light and dark cycles. The model illustrates the relationship between light/dark conditions, Drp1 levels and mitochondrial recruitment, and the resulting normal versus enhanced priming states.

Therefore, we asked if such a mitochondrial priming of GSC proliferation for dietary supplement stimulated egg production sustains in the Cry mutants where the light-dark rhythm is perturbed. Interestingly, stimulated Cry^out^-GSCs and their progenies in LD exhibit a pronounced decline in mitosis (rate or length) compared to the W^1118^-GSCs (Fig. 4D-E). Consistently, both Cry^b^ and Cry^out^ mutants fail to exhibit the stimulated egg production in LD, LL or DD, in a graded manner between the mutants, with no consistent difference in the basal levels (Fig. 4F). These data suggest defective priming of the ovarian stem cells in the Cry mutants. Moreover, the following attributes of entrainment of egg production indirectly indicate interaction of protein-rich supplement with both light-dependent and light-independent actions of Cry in modulating ovarian stem cells (Fig. S4E): **a)** inhibitory impact of Cry mutants on egg production progressively increases over 10 days; **b)** basal free running rhythm (DD) is established in W^1118^-GSCs after 7 days, which is abrogated after stimulation; **c)** total loss of CRY (Cry^out^ mutant) maintains a continuous basal rhythm only in LD, whereas stimulation establishes it beyond 7 days; **d)** light insensitivity (Cry^b^ mutant) maintains the free running basal rhythm before 7 days and its gradual dampening is prevented with stimulation. Stimulation of egg production remains inhibited in the Cry^b^ and Cry^out^ mutants in the HSD environment that supports self-renewal/proliferation of the ovarian stem cells (Fig. S4F, Fig. S3E).

To identify if any particular amino acid in a protein-rich supplement can drive the mitochondrial priming of GSCs to boost egg production, we screened all essential amino acids. Yeast extract served as a positive control during the screen, whereas the holidic diet established the benchmark concentration for individual amino acid supplementation (see Methods). We evaluated a spectrum of doses for each amino acid, ranging from 100-fold below to 10-fold above holidic diet levels, applying these to our basal diet to calculate supplement concentrations. Threonine stimulates egg production to the control level over 12 days of egg development, with no stimulation by direct nutritional influence over 2 days unlike Phenylalanine and Valine (Fig. 4G); greater than optimal amino acid supplementation lowers egg production or even causes organismal death (Fig. 4H, Fig. S4F), likely due to amino acid imbalance (Grandison et al. 2009; Knopf and Lamming 2026). Importantly, Threonine supplementation alone elevates mitosis (rate or length) of the W^1118^-GSCs to the level of total protein-rich supplement, which does not happen with Phenylalanine (Fig. 4H). Such Threonine driven stimulation of GSC mitosis involves Cry action since Threonine suppresses basal mitosis in Cry^out^-GSCs and their progenies (Fig. 4H); no impact on organismal survival is detected. Phenylalanine sustains the same level of mitosis of Cry^out^-GSCs and its progenies as total protein-rich supplementation, both being pronouncedly lower than W^1118^-GSCs.

To confirm if Threonine driven mitosis boost in GSCs requires them to be primed in a Drp1-Opa1 and light-dark cycle dependent manner, we quantified mitosis in Drp1 knockdown (primed) or Opa1 knockdown (defective priming) GSCs and their progenies in mid-light and mid-dark (Fig. 4I, J). Interestingly, overall protein-rich supplementation stimulates mitosis (rate or length) more in the dark in control. Whereas, mitosis boost by Threonine and Drp1-knockdown, and suppression by Opa1-knockdown, happens in both mid-light and mid-dark. Moreover, Threonine dramatically suppresses mitosis in the primed Drp1 knockdown cells below the basal and significantly rescues mitosis defect in Opa1-knockdown in both mid-light and mid-dark. Given this was not observed with Phenylalanine (Fig. 4I), the data demonstrates that the specific impact of Threonine on mitosis requires higher levels of Drp1 and its action is downstream to Opa1. Whereas, Phenylalanine suppresses basal mitosis of the control and Opa1 knockdown cells.

In summary, protein-rich supplement dependent mitochondrial priming of W^1118^-GSCs requires cycling through light and dark, while stimulated mitosis happens predominantly in dark to support egg production. Thus, absence of Cry functionality prevents GSC priming and disables them from responding to protein-rich supplements to stimulate egg production. On the other hand, the stimulated mitosis of Drp1-knockdown GSCs happens additionally in light cycles leading to enhanced egg production. Similarly, the essential amino acid Threonine alone can drive the Cry and Drp1 dependent priming of GSCs by additionally boosting mitosis in the light cycle that activates Cry and has higher level of Drp1 (Fig. 3A-C). Thus, Threonine rescues the defective priming in Opa1 deficient GSCs.

## Discussion

Although various mitochondrial functions regulate stem cell activation, self-renewal/proliferation, differentiation (Y. Wang et al. 2024; Chakrabarty and Chandel 2021), the prevalence and regulation of the proposed ‘primed’ stemness state with distinct mitochondrial properties remains elusive. Scattered literature indicates a high potency stemness state that has characteristic mitochondrial membrane potential, content and fission-fusion dynamics, as detected in various lineages across various organisms (Khacho et al. 2016; Meacham et al. 2022; Charmpilas and Tavernarakis 2020; Deng et al. 2018; Seo et al. 2023). Using an in vitro cell system, we previously reported that mitochondrial priming of stemness to support neoplasticity can be achieved by tuning mitochondrial fission protein, Drp1, to a ‘goldilocks’ level (Spurlock et al. 2021b). Here, we demonstrate that in vivo tuning of Drp1 to an optimal level sustains a mitochondrial primed state of higher self-renewal/proliferation and differentiation in *Drosophila* ovarian stem cells towards boosting egg production. Our quantitative data from gene-environment interaction studies and near super resolution microscopy is consistent with the following mechanistic model of mitochondrial priming of *Drosophila* ovarian GSCs by regulated repression and activation of Drp1 (Fig. 4K):

In the normal dark cycle of priming with reduced Drp1 protein levels, protein-rich supplement stimulates Drp1 recruitment onto mitochondrial-units in a characteristic number of small-immature clusters to maintain restricted mitochondrial fission and thus allow unopposed mitochondrial fusion (likely Opa1 driven) (Fig. 2, 3). In the light cycle with higher Drp1 levels, protein-rich supplement stimulates the Drp1 clusters to mature to large Drp1-clusters to activate mitochondrial fission (Fig. 2, 3). Such a cycling of Drp1 activity allows elevated mitosis in self-renewing GSCs and their immediate differentiated progenies primarily in the dark cycle to support egg production (Fig. 1, 4). In the absence of Cry functionality, Drp1 levels are constitutively high and Drp1 is recruited aberrantly to mitochondria in the light-cycle while its characteristic mitochondrial recruitment of the dark-cycle is perturbed (Fig. 3). This disables mitochondrial priming for protein-rich supplement driven boost in egg production when Cry functionality is absent (Fig. 4). Constitutive mild Drp1 repression driven enhanced priming allows protein-rich supplement stimulated characteristic mitochondrial Drp1 recruitment to elevate mitosis in both light and dark cycles (Fig. 2, 4). This elevates overall self-renewal and differentiation of the GSCs to boost egg production in an Opa1 dependent manner (Fig. 1, 4). Such a mitochondrial priming of GSCs is supported by the essential amino acid Threonine alone in a Cry dependent manner, which elevates mitosis in light in a Drp1 dependent manner and rescues defective mitosis in Opa1 deficient GSCs (Fig. 4).

Our data demonstrates that priming of GSCs is inhibited in Opa1 deficient cells (Fig. 1F-K, Fig. 4I-J), and potentially in Marf-1 deficient cells (not shown). Threonine driven rescue of mitosis in Opa1 deficient GSCs and suppression of it in Drp1 deficient GSCs indicate that Threonine action is downstream of Opa1 and still needs upstream Drp1 action (Fig. 4I). Threonine uniquely supports certain stem cells through mitochondrial one-carbon metabolism in addition to supporting TCA cycle driven mitochondrial energetics (Sahoo et al. 2021; J. Wang et al. 2009; Van Winkle and Ryznar 2019). Therefore, our data points to the following testable hypothesis on the mechanism of Threonine driven boost of mitochondrial priming of stem cells: Drp1 driven mitochondrial recruitment to the large mitochondrial units sustained by Opa1 (and Marf-1) driven mitochondrial fusion support Threonine driven one carbon metabolism and mitochondrial bioenergetics for boosting GSC mitosis and self-renewal both in the light and dark cycles (Fig. 4K).

The observation that quiescent GSCs appear to reside in G2-M (Fig. S1G) and their Drp1 dependent mitochondrial priming involves modulation of mitosis (Fig. 1I, K Fig. 4C,I,J) raises the possibility that the regulated repression and activation of Drp1 happens through G2-M regulation. Given Drp1 integration into cell cycle regulation includes its activation during normal mitosis (Spurlock et al. 2020; Chen and Chan 2017), mild Drp1 repression is indeed expected to slow down mitosis and thus increase the length of mitosis in dark (Fig. 4K). The G2-M stage of cell cycle state is a key determinant of stemness properties (Van Oudenhove et al. 2016; Garyn et al. 2024; Soufi and Dalton 2016), while mitochondrial redox and metabolic regulatory properties are key for active cell cycle modulation (Y. Wang et al. 2024; Kirova et al. 2022). Indeed, regulation of adult cell cycle in stem cells (and others) is intimately connected to Cry dependent light-dark cycles (Gliech and Holland 2026; Brown 2014). While Cry can function in a light independent manner (Griffin et al. 1999; Lamia et al. 2011; Zhang et al. 2010), non-canonical direct involvement of Cry has been shown in pluripotent stem cells (Dierickx et al. 2018; Sato et al. 2023). Our confirmation of active Cry promoter in ovarian GSC and FSC lineages (Fig. S3F-G) and opposite phenotype between W^1118^ and Cry^out^ GSCs indicate Cry action may prevent the characteristic mitochondrial Drp1 recruitment and organization in light cycle and allow it in dark only when stimulated. This conception opens the question of whether the demonstrated Cry involvement involves combination of light sensitive and insensitive non-canonical action of Cry (Fig. 4K). These non-canonical functions of Cry involvement in the peripheral tissues including stem cells, possibly couples the temporal signal to local metabolic state to influence the stem cell proliferation or fate. We, along with others (Andersen et al. 2023; Tulina et al. 2014) have demonstrated that mitosis (rate / length) is preferentially increased during the dark relative to the light phase (Fig. 4J). Nonetheless, lineage specific differences may exist in the molecular circuitry of connectivity between mitochondrial priming, cell cycle and light-dark cycle, as highlighted by the difference in priming levels of Drp1 and cell cycle state between GSCs and FSCs (Fig. 1B,L, S1G).

The disease relevance of mitochondrial priming of *Drosophila* ovarian stem cells may potentially extend to diabetes, cancer and aging, where adult stem cell functionalities are affected. Indeed, our data demonstrates that mitochondrial priming of *Drosophila* ovarian germline and somatic stem cells can remarkably overcome the inhibition of the high sugar dietary environment and can even sustain elevated stem cell proliferation in that environment. Therefore, our study revealing and explaining the integration of light and dietary cues to regulate adult stem cells via precise mitochondrial regulation hold implications in the areas of regenerative health and disease etiology and management.

## Materials and Methods

### *Drosophila* maintenance and crosses

*Drosophila* strains were maintained on standard cornmeal medium under a 12:12 light: dark (LD) cycle at 25°C, unless otherwise specified. All germline RNAi-mediated knockdowns were driven by nanos-GAL4 or nanos-GAL4 combined with UAS-Dicer-2. The somatic knockdown was driven by Gal109-30. These driver lines were crossed with UAS-RNAi lines targeting genes such as Drp1 and Opa1. All stocks are listed in the Key Resources table. Following the crosses, progeny carrying the appropriate GAL4/UAS combinations were identified based on the relevant genetic markers and selected for downstream experiments.

To visualize expression of Cryptochrome promoter, a Cry-Gal4 driver line was combined with a UAS-RFP reporter line.

### Dietary manipulation

All fly stocks were maintained in standard cornmeal medium, which was prepared at a final volume of 1L using the following components: 9 g agar, 51 g maize flour, 51 g sugar and 18 g yeast. Methyl paraben (3 g in 6 mL ethanol) and propionic acid (3 mL) were added after cooling the medium to approximately 55°C. For yeast-stimulated diet, granular yeast was added to the food vials. For dietary manipulations, flies were provided either with a control diet or a high sugar diet (six-fold increase in sucrose content), with or without additional yeast supplementation (Palanker Musselman et al. 2011). For amino acid supplementation experiments, holidic diet (Piper et al. 2014) was used as a benchmark to determine concentrations. We determined the additional supplemental concentration required to elevate our basal diet (estimated from the raw ingredients) to holidic diet for each essential amino acid. A systematic screening was then performed using a range of doses, including 100-fold lower and 10-fold higher concentrations to the estimated supplemental concentration, to identify the most effective dose. Each amino acid was then tested by addition of these 3 calculated concentrations to the basal food to determine the specific dosage that maximizes egg-laying and egg-production.

### Egg production assay

The physiological outcome of stem cell functionality was assessed using egg production assays as a readout under various conditions (Drummond-Barbosa and Spradling 2001; Templeman and Murphy 2018). Post-eclosion mated females (aged 1-5 days) were initially housed in vials with food dependent on the specific assessment. Depending on the assessment, the flies were subjected to their respective light dark regimes - LD (12:12 light: dark), LL (24 hrs. light), and DD (24 hrs. dark) at 25°C. The flies were fed either basal cornmeal, stimulated (with granulated yeast), control diet, high sugar diet or amino acid supplemented food with their respective concentration. 4 males and 4 females were then transferred to individual conical set-ups 2 days before the reading with food plates. The food plates were provided with their respective diet along with a few drops of food colour to easily visualize the eggs.

Laid eggs were manually counted at the end of the light and dark phases on specified days (e.g. Day 2 and Day 12) under different light and dietary regimens. The values at the end of light and dark phases were summed to get the total egg count per day.

### Immunohistochemistry and Immunoblotting

Immunohistochemistry was done following our optimized protocol (Parker et al. 2017). In brief, *Drosophila* ovaries were dissected out in RT Grace’s insect medium and fixed in 4% freshly prepared paraformaldehyde for 15 minutes. The samples were permeabilized in PBS containing 0.8% Triton X-100 (PBST), blocked in PBST containing 2% BSA and followed with primary antibody incubation (anti-DRP1 (custom-generated and validated), anti-ATPB (mitochondrial marker – Cat# Ab14730), anti-p-HistoneH3 (mitotic marker – Cat# sc-374669)) for 3 h in RT or overnight at 4°C. The samples were washed with PBST thrice for 15 min each. Samples were then incubated with Alexa Fluor 488, Cy3, or Cy5-conjugated secondary antibodies (Jackson ImmunoResearch Laboratories) in PBST for 1 h, followed by a 30-minute counterstaining of nuclei using Hoechst after 2 washes in PBST of 15 min each.

For immunoblotting, protein lysates were prepared from 40-45 ovaries, dissected in RT Grace’s insect medium. The anterior region of each ovarioles (germarium and stages 2-3) was further dissected out and immediately transferred into 100 μL of ice-cold RIPA buffer (50mM Tris HCl pH 8, 0.1% SDS, 150mM NaCl, 1% Triton X-100, 10mM Na3VO4, 50mM NaF, 2mM PMSF in Ethanol and 1:100 Protease inhibitor cocktail). Samples were immediately flash-frozen in liquid nitrogen and stored at –80 °C until further processing. For lysis, frozen samples were thawed on ice, homogenized using a handheld micro-pestle and further homogenized by passing through a syringe (5 passes). Lysates were incubated on ice for 15 minutes and centrifuged at 14,000 rpm for 10 minutes at 4 °C. The supernatant was collected and protein concentration was estimated using the BCA assay.

Protein samples (100 μg per lane) were boiled in Laemmli buffer at 95 °C for 5 minutes and resolved on SDS-PAGE gels. Proteins were transferred onto PVDF membranes and probed using the following primary antibodies after blocking for 1 hour in 5% BSA - anti-DRP1, anti-ATPB, and anti-β-Actin (Cat# sc-47778) as loading control. Compatible HRP-conjugated secondary antibodies were utilised, followed by detection using chemiluminescence. Densitometric analysis of protein bands was performed using ImageJ, with normalization by loading controls, as appropriate.

### Confocal microscopy

All microscopy was performed using a Zeiss LSM 900 equipped with an Airyscan2 super-resolution module. The microscope was configured with a 63X Plan-Apochromat 1.4 NA oil immersion objective, which was utilised for all acquisitions. Appropriate excitation laser lines were used for each fluorophore or secondary antibody used. Three-dimensional image stacks were acquired using optical zoom (2X), at 1 Airy unit pinhole, and 0.5 µm Z-interval for all lineage tracing, pH3 analysis and average intensity analysis. Whereas, to trace the developmental outcome of GSC and FSC lineages across the entire ovariole chain, scans were conducted at a 0.5X optical zoom setting. Same acquisition settings were maintained across all images of a particular experiment to facilitate comparison between them.

Quantification of average signal levels for Fly-FUCCI, AtpB, and Drp1 within the germarium was conducted utilizing manually defined regions of interest (ROIs) in Zen Blue Software incorporating appropriate background correction measures.

For the Mitochondrial and Drp1 cluster analysis pipeline, the anterior region of the germarium was scanned using the AiryScan2 module with the 63X objective. Here, 16-bit 3D image stacks were acquired using optical zoom (5X), at 1 Airy unit pinhole, and 0.14 µm Z-interval. The images were ‘AiryScan processed’ using Zeiss Zen Blue Software.

### Phospo-Histone H3 quantification

Phospho-histone H3 (pH3) positive cells were quantified from germaria of dissected and appropriately stained ovarioles. All germaria with or without pH3 signal were considered. The number of pH3 positive nuclei counted in Region 1 (anterior tip excluding cap cells to the boundary where FSCs emerge) were aggregated across all germaria and expressed as a percentage of pH3 positive cells.

### Clonal Strategy and Analysis

Homozygous drp1 mutant clones were generated by FLP-FRT mediated site-specific mitotic recombination in the background of heterozygous tissue (ref). Drp1-mutant or control FRT40A clones were generated by crossing *hsflp ubiGFP/CyO* males with virgin females of *FRT 40A/CyO*, *drp1^2^ FRT 40A/CyO*, *drp1^1^ FRT 40A/CyO* or *drp1^KG03815^ FRT40A/CyO* alleles. Clones in the desired progeny were induced using heat shock at 37.5°C (1 hour, twice a day with 6 hour gap) for 2 consecutive days. Following this, they were transferred into their respective dietary and light regimes and maintained for 12 days. This allows the egg chambers with transient clones clear out as eggs laid, leaving only the fixed stem cell clones for lineage tracing analyses.

The GFP negative clonal GSCs self-renew, proliferate and differentiate to form the progressively growing GFP negative germ line cysts at stages of egg development. Each GFP negative clonal FSC differentiate into GFP negative FCs and contributes equally to the egg chamber. Therefore, for every ovariole chain, measurements were conducted in the germarium and in their traced lineages reflected in their egg chambers. Ovarioles were segregated into 2 categories, those with Germline clone (GSC/cyst) and those with Somatic clone (FSC/FC). Subsequent analyses were restricted exclusively to ovarioles consisting of either type of clone. The GSC/ FSC properties quantified, as described below, are described in Fig. S1B.

Abundance of total clonal GSC/FSC normalized to the total number of GSCs or FSCs (2 per germarium) in the group indicates overall clonal GSC/FSC abundance. Of these, germarium was further categorised into 0,1, or 2 Clonal GSCs. Abundance of germarium with 0 clonal GSCs but with at least one clonal germline cyst indicates GSC loss. Given a clonal GSC would self-renew/proliferate to form the 2nd clonal GSC, the ratio of abundance of germarium with 2 vs 1 clonal GSC indicates GSC self-renewal/proliferation. Abundance of clonal cysts in a given ovariole chain normalized to the total number of cysts identified in that chain indicates GSC differentiation. The longitudinal axis length of a clonal cyst indicates germline cyst size.

The abundance of germarium with 0 clonal FSCs but with 1 or more FC clones in the ovariole chain indicates FSC loss. Of the germarium with 1 or 2 clonal FSCs, the number of FC clones in each egg chamber normalized to the number of clonal FSC in its germarium indicated the number of FSC divisions. This reflects the FSC Self renewal/proliferation with successful differentiation. Given the FSC would divide only once per cyst that contributes equally to the FCs encapsulating the germline, a division more than once would be reflected in the log2 values of the abundance of FCs. Those that divide greater than once per ovariole chamber will be reflected as >0 value in the log2 transformed value.

### Connected-component analysis to characterize Mitochondrial and Drp1 clusters

This image analysis pipeline was created combining tools from Zeiss Zen Blue , Cell Profiler 4.2.8 and MATLAB R2025a (Fig. S2A). The major steps are described below:

**Step 1**: a. Deconvolution: This was performed on Airyscan-processed images using the “Fast Iterative algorithm, Good quality” setting.

**Step 1:** b. Cell Isolation: GSCs were identified as the two distinct large cells at the anterior tip of each germarium, either by their nuclei or by the absence of GFP for lineage tracing analyses. GSC - stack inputs were generated by drawing ROI around each GSC, and the cells were cropped and saved with unique Cell-IDs. Each GSC Z-stack was separated into discrete individual Z-slice frames and organized into separate mitochondrial and Drp1 folders within the corresponding Cell-ID directory. Images were named systematically as Cell-ID_Mitochondria_Z-Slice# for the and Cell-ID_Drp1_Z-Slice#.

**Step 2: a. Thresholding:** The individual Z-Slices were processed using CellProfiler. ‘Primary objects’ were identified in each Z-slice using the ‘IdentifyPrimaryObjects’ module with “Adaptive” thresholding strategy and “Otsu” thresholding method. For pipeline optimization, multiple “typical diameter of objects” ranges were tested. Then diameter was set to 10–40 pixels for mitochondrial objects (default settings) and 5–30 pixels for Drp1 clusters. Diameters below the selected minimum resulted in the inclusion of background pixels, whereas increasing the maximum diameter led to the merging of adjacent objects.

**Step 2: b. Binarization:** The ‘ConvertObjectsToImage’ and ‘SaveImages’ modules were used to generate binary images of the thresholded images for individual channels, which were saved as CellID_Mitochondria_Mask_ZSlice# and CellID_Drp1_Mask_ZSlice#, respectively.

**Step3: 3D layering and rendering:** The individual Z-slice binary masks were input into MATLAB and were reconstructed into 3D volumes by sequentially stacking. The reconstructed volumes were visualized using MATLAB’s “volshow” function.

**Step 4: Drp1 filtering:** The Drp1 pixels were first identified and categorized into mitochondrial and non-mitochondrial pools based on their direct X, Y, and Z coordinate overlap.

**Step 5: a. Connected component analyses:** The mitochondrial pool of Drp1 pixels and all Mitochondrial pixels were subjected to cluster detection. To perform this, we applied connected component analysis to the 3D binary reconstructions via the MATLAB “bwconncomp” function, with “6-connectivity”. This method assigned unique ClusterIDs to each identified structure, while the total pixel count for each cluster was its size. Clusters ≤5 pixels in size were excluded from further analysis, and the remaining clusters were reassigned to unique Cluster IDs. This threshold was established based on the resolution limit of the microscope and the individual pixel size to minimize the inclusion of potentially artifactual clusters. The individual mitochondrial-units with their overlapping Drp1 clusters were visualized using the “volshow” function on MATLAB.

**Step 5: b. Data extraction:** All mitochondrial-units and Drp1 cluster metrics, including Cluster IDs, cluster sizes and the X, Y, and Z coordinates of every pixel, were recorded in a comprehensive Excel spreadsheet and saved within the corresponding Cell-ID directory.

**Step 6: Feature computation:** Utilizing this dataset we looked at various metrics to understand the mitochondrial and Drp1 properties at the Cell-level and individual mitochondrial- unit and Drp1 cluster level. The properties primarily studied were -

- Number of Drp1 pixels in each GSC (% fraction) -Percentage of the total Drp1 pixels identified within each GSC that directly overlap with mitochondria, representing the mitochondria-associated Drp1 pool, versus those that do not overlap with mitochondria, representing the non-mitochondrial Drp1 pool.
- Fraction overlap of Mito-Drp1 pixels in each GSC (%) - Percentage of mitochondrial pixels that overlap with Drp1 pixels in each GSC
- Number of mitochondrial Drp1-clusters in each GSC - Total number of Drp1 clusters (after Drp1 filtering described in step 4) in each GSC
- Average size of mitochondrial Drp1-clusters in each GSC - Average size of the Drp1 clusters in each GSC
- Number of mitochondria-units in each GSC - Total number of mitochondrial-units in each GSC
- Average Size of mitochondrial-units in each GSC - Average size of the mitochondrial-units in each GSC
- Number of Drp1-clusters on each mitochondrial-unit -The total number of Drp1 clusters associated with each mitochondrial unit.
- Average number of Drp1 clusters on each mitochondrial-unit in each GSC - Average number of Drp1 clusters associated with each mitochondrial-unit within an individual GSC.
- Ln (Mitochondrial-unit size) - Log transformed size of individual Mitochondrial-unit
- Ln(Size of Drp1-cluster on each mitochondrial-unit) - Log transformed size of Individual Drp1-clusters on each Mitochondrial-unit

### Statistical Analysis

All plots and statistical analyses were performed using R (version 4.4.1) using R studio. All statistical comparisons between groups were performed using the non-parametric Mann–Whitney U test. Fisher’s exact test was used to assess differences in clonal abundance in the dot plots. P-values less than 0.05 were considered statistically significant and denoted as follows: p< 0.05 *, p< 0.01 ** and p< 0.001 ***.

## Acknowledgements

The research was supported by DBT-Wellcome Trust India Alliance (IA/S/20/2/505198/WTDBT); YS was supported by Ashoka University.

## Key Resources Table

| REAGENT/RESOURCE | SOURCE | IDENTIFIER |
| --- | --- | --- |
| **Antibodies** |  |  |
| Rabbit anti-DRP1 | Boster Bio | This paper; Cat#: DZ41314 |
| Mouse anti-ATP-B | Abcam | Cat#: Ab14730 |
| Mouse anti-p-Histone H3 | Santa Cruz Biotechnology | Cat#: sc-374669 |
| Mouse anti-β-Actin | Santa Cruz Biotechnology | Cat#: sc-47778 |
| Rabbit anti-IgG- HRP Linked | Cell Signaling Technology | Cat#: 7074P2 |
| Mouse anti-IgG- HRP Linked | Cell Signaling Technology | Cat#: 7076P2 |
| Cy<sup>TM</sup>3 AffiniPure® Goat Anti-Mouse IgG (H+L) | Jackson Immuno research | Cat#: 115-165-146 |
| Cy<sup>TM</sup>3 AffiniPure® Goat Anti-Rabbit IgG (H+L) | Jackson Immuno research | Cat#: 111-165-144 |
| Cy<sup>TM</sup>5 AffiniPure® Goat Anti-Mouse IgG (H+L) | Jackson Immuno research | Cat#: 115-175-146 |
| Alexa Fluor® 488 AffiniPure® Goat Anti-Rabbit IgG (H+L) | Jackson Immuno research | Cat#: 111-545-144 |
| **Chemicals** |  |  |
| Grace's Media (Insect Dissecting Medium) | Sigma-Aldrich | Cat#: G9771 |
| Paraformaldehyde (EM grade) | Electron Microscopy Solutions | Cat#: 15711 |
| Triton X-100 | SRL | Cat#: 64518 (2024271) |
| Sodium chloride | Sigma-Aldrich | Cat#: S9888 |
| Potassium chloride | Sigma-Aldrich | Cat#: P3911 |
| Sodium phosphate dibasic | Sigma-Aldrich | Cat#: S9763 |
| Potassium phosphate monobasic | Sigma-Aldrich | Cat#: P0662 |
| BSA | Sigma-Aldrich | Cat#: A9647 |
| Hoechst-33342 | Invitrogen | Cat#: H3570 |
| Fluoromount G | Southern Biotech | Cat#: 0100-01 |
| Nail polish | MAC | Amazon |
| Sodium Lauryl Sulphate (SDS, SLS) | SRL | Cat#: 54468 |
| Tris Buffer AR, ACS | SRL | Cat#: 71033 |
| Tris HCl | SRL | Cat#: 99438 |
| Glycine | SRL | Cat#: 64072 |
| Sodium Orthovanadate | SRL | Cat#: 86766 |
| Sodium Fluoride | SRL | Cat#: 29821 |
| Phenylmethane Sulphonyl Fluoride | SRL | Cat#: 87606 |
| Magnesium Sulphate | Sigma Aldrich | Cat# - M7506 |
| Calcium Chloride dihydrate | Sigma Aldrich | Cat# - C5080 |
| Pierce Protease and phosphatase inhibitor | Thermo Scientific | Cat#: A32959 |
| BCA Protein estimation Kit | Merck Millipore | Cat#: 71285-3 |
| 2X Laemmli Buffer | BIORAD | Cat#: 1610737 |
| Ammonium persulphate | Fisher Scientific | Cat#: BP179 |
| TEMED | Merck Millipore | Cat#: T9281 |
| Tween-20 | Promega | Cat#: H5152 |
| Methanol | Spectrochem | Cat#: 71301 |
| EMSURE Ethanol | Sigma Aldrich | Cas#: 64-17-5 |
| 30% bis acrylamide 29:1 | BIORAD | Cat#: 1610156 |
| Clarity Western ECL Substrate | BIORAD | Cat#: 1705061 |
| Ponceau S Solution | Sigma Aldrich | Cat#: P7170 |
| Precision Plus Protein Dual Color Standards | BIORAD | Cat#: 1610374 |
| Agar | SRL | Cat#: 19661 |
| Methyl Paraben | Sigma Aldrich | Cat# 61862805001730 |
| Propionic Acid | SRL | Cat# 12931 |
| Sugar (Sulphur less) |  | Amazon |
| Maize Fluor |  | Amazon |
| Yeast |  | Amazon |
| Sucrose | Sigma Aldrich | Cat# S0389 |
| Yeast Extract | Himedia | Cat# RM027 |
| Peptone | SRL | Cat# 95292 |
| **Experimental models: *Drosophila melanogaster* strains** |  |  |
| *w*1118 | (Das et al. 2024) | Gift; BDSC_5905 |
| *Canton S* |  | BDSC_64349 |
| *w*[1118]; *P*{*ry*[+*t*7.2]=*neoFRT*}40A/CyO;<br>*P*{*ry*[+*t*7.2]=*ey-FLP.N*}6, *ry*[506] | (Mitra et al. 2012) | BDSC_8212 |
| *drp1*<sup>KG03815</sup> *FRT* 40A/CyO | (Mitra et al. 2012) | BDSC_13510 |
| *y*[*d2*] *w*[1118] *P*{*ry*[+*t*7.2]=*ey-FLP.N*}2 *P*{5xg\|BS-lacZ.38-1}TPN1; *P*{*y*[+*t*7.7] *ry*[+*t*7.2]=*Car20y*}25F<br>*Drp1*[1] *P*{*ry*[+*t*7.2]=*neoFRT*}40A/CyO,<br>*P*{*w*[+*mC*]=*GAL4-Kr.C*}DC3, *P*{*w*[+*mC*]=*UAS-GFP.S65T*}DC7 | (Verstreken et al. 2005) | Gift; BDSC_24885 |
| *y*[*d2*] *w*[1118] *P*{*ry*[+*t*7.2]=*ey-FLP.N*}2 *P*{5xg\|BS-lacZ.38-1}TPN1; *P*{*y*[+*t*7.7] *ry*[+*t*7.2]=*Car20y*}25F<br>*Drp1*[2] *P*{*ry*[+*t*7.2]=*neoFRT*}40A/CyO,<br>*P*{*w*[+*mC*]=*GAL4-Kr.C*}DC3, *P*{*w*[+*mC*]=*UAS-GFP.S65T*}DC7 | (Verstreken et al. 2005) | Gift; BDSC_24899 |
| *w*[1118]; *P*{*w*[+*mC*]=*Ubi-GFP(S65T)nls*}2L<br>*P*{*ry*[+*t*7.2]=*neoFRT*}40A/CyO | (Mitra et al. 2012) | BDSC_5629 |
| *y*[1] *w*[*]; *P*{*w*[+*mC*]=*GAL4-nanos.NGT*}40 | (Chowdhary et al. 2020) | Gift; BDSC_4442 |
| *P*{*w*[+*mC*]=*UAS-Dcr-2.D*}1, *w*[1118];<br>*P*{*w*[+*mC*]=*GAL4-nanos.NGT*}40 | (Ni et al. 2008) | BDSC_25751 |
| *y*[1] *w*[*]; *P*{*w*[+*mW.hs*]=*GawB*}109-30/CyO | (Hartman et al. 2010) | BDSC_7023 |
| *y*[1] *sc*[*] *v*[1] *sev*[21]; *P*{*y*[+*t*7.7]<br>*v*[+*t*1.8]=*VALIUM20-mCherry*}attP2 | (Ni et al. 2008) | Gift; BDSC_35785 |
| *w*[*]; *Bl*[1]/CyO; *P*{*w*[+*mC*]=*UAS-Opa1.miRNA.CDS*}3 | (Uttekar et al. 2024) | Gift; BDSC_67159 |
| *y*[1] *sc*[*] *v*[1] *sev*[21]; *P*{*y*[+*t*7.7]<br>*v*[+*t*1.8]=*TRiP.HMS00349*}attP2 | (Uttekar et al. 2024) | Gift; BDSC_32358 |
| *w*1118; *P*{*GD10456*}v44156/TM3 |  | Gift; VDRC_GD-44156 |
| *w[\*]; P{w[+mC]=UAS-Drp1.RNAi.CDS.S}3/T(2;3)TSTL, CyO: TM6B, Tb[1]* |  | Gift; BDSC_67160 |
| *w[1118]; ss[1] cry[b]* | (Stanewsky et al. 1998) | BDSC_80921 |
| *w[1118]; P{w[+mC]=XP}cry[d10630]* | (Yoshii et al. 2008) | BDSC_19331 |
| *w[1118]; Kr[lf-1]/CyO, P{ry[+t7.2]=en1}wg[en11]; P{w[+mC]=Ubi-GFP.E2f1.1-230}5 P{w[+mC]=Ubi-mRFP1.CycB.1-266}4/TM6B, Tb[1]* | (Zielke et al. 2014) | BDSC_55099 |
| **Software and algorithms** |  |  |
| ImageJ 1.53t (FIJI) | (Schindelin et al. 2012) | https://imagej.nih.gov/ij/index.html |
| R Version 4.3.3 | R Core Team (2024) | https://cran.rstudio.com/ |
| MATLAB | The MathWorks, Inc. | https://in.mathworks.com/products/matlab.html |
| CellProfiler 4.2.8 | (Carpenter et al. 2006) | https://cellprofiler.org/ |
| **Other** |  |  |
| LSM 900-Confocal Microscope with AiryScan2 | Carl Zeiss |  |
| Dissecting Microscope | Carl Zeiss | Stemi 305 |
| Serological pipettes | Corning | Cat#: 4487 |
| Tissue roll | - | Amazon |
| Auto-pipette tips | Fisher Scientific | - |
| Falcons, 15 mL and 50 mL | Corning | Cat#: 430055, 430829 |
| Glass slides | Azer Scientific | EMS200B |
| Coverslips | Blue Star | 22mmX22mm |
| Micro-pestles | Praveen Scientific | Cat#: 23251 |
| Syringe with needle | Dispovan | Cat#: 13481 |
| Mini-PROTEAN® Short Plates | BIORAD | Cat#: 1653308 |
| PVDF Membrane | BIORAD | Cat#: 1620177 |
| Mini-PROTEAN® Spacer Plates with 1.5 mm Integrated Spacers | BIORAD | Cat#: 1653312 |
| Autoclave | Tomy-SX-700 | - |
| Shaker rocker | Rocking shaker II, BOEKL | - |
| Class-II bio-safety hood with laminar flow | Microfilt | - |
| Auto-pipettes | Eppendorf | - |
| Incubator | PHCBI | Cat#: MIR-554 |
| Pipette gun | Drummond Scientific | - |
| Dumont #5 Forceps | Fine Science Technologies | 11251-20 |
| MultiDish 8 Well Square, 24x33mm wells | Electron Microscopy Sciences | 70443 |
| Fly Vials | Laxbro | BT-30A(25mmX90mm) |
| Fly Bottles | Laxbro | FLBT-20 (Item Code: 110451 / 110452) |
| Fly Conicals | Fisher Scientific | AS-355 |
| Fly Flugs Vials | Fisher Scientific | AS-273 |
| Fly Flugs Conical | Fisher Scientific | AS-277 |

## Supplementary Figure Legends

**Supplementary Fig 1.**
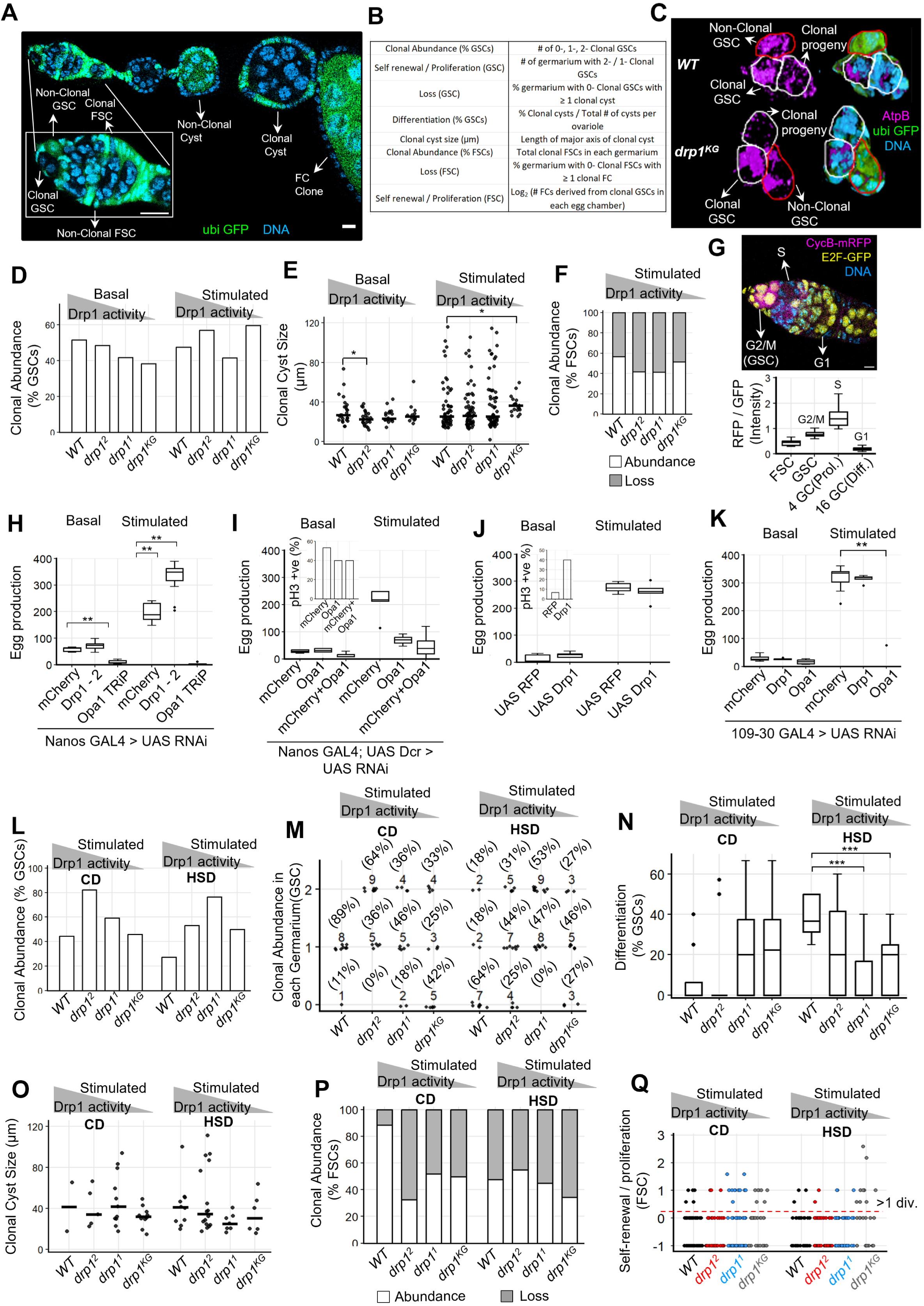
A) Representative confocal optical slice indicating a clonal ovariole with clonal GSC, FSC, Cyst and FCs stained with Hoechst DNA stain ; scale bar : 5 μm B) Table describing clonal parameters measured in lineage tracing analyses shown in Fig. 1. C) A section of 3D rendered micrograph of GSC and its immediate progeny stained for AtpB and Hoechst DNA stain in WT and *drp1^KG^* germarium. D) Bar plots indicating clonal GSC abundance (%) (N(germaria)-Basal: WT-33, *drp1^2^*-33, *drp1^1^*- 24, *drp1^KG^*-17; N(germaria)-Stimulated: WT-41, *drp1^2^*-32, *drp1^1^*-26, *drp1^KG^*-26). E) Dot plots depicting clonal cyst sizes (μm) (N(cyst)-Basal: WT-25, *drp1^2^*-23, *drp1^1^*-18, *drp1^KG^*- 12; N(cyst)-Stimulated: WT-56, *drp1^2^*-59, *drp1^1^*-64, *drp1^KG^*-23). F) Stacked bar plots indicating clonal FSC abundance(black) and Loss(grey) (%) (N(germaria)- Stimulated: WT-34, *drp1^2^*-26, *drp1^1^*-32, *drp1^KG^*-33) G) Representative confocal optical slice of germarium of Fly-FUCCI line (above), with the box plot (below) showing quantification of the RFP / GFP intensity ratio indicating cell cycle stages. (N : 29 germaria) H) Boxplots showing egg-production on Day 12 following germline-specific knockdown of mCherry, Drp1, or Opa1. (N-Basal: mCherry RNAi-5, Drp1 RNAi-13, Opa1 RNAi-8; N-Stimulated: mCherry RNAi-13, Drp1 RNAi-13, Opa1 RNAi-8) I) Boxplots showing egg-production on Day 12 following germline-specific knockdown of mCherry, Opa1 and simultaneous knockdown of mCherry and Opa1 (N:5/genotype/condition). Inset shows abundance of pH3-positive cells (%) in Region 1 of germaria (N : 15 germaria for each genotype). J) Boxplots showing egg-production on Day 12 following germline-specific over-expression of RFP and Drp1 (N : 5/genotype/condition). Inset shows bar plot of % pH3-positive nuclei in Region 1 of germaria (N : 15 germaria for each genotype). K) Boxplots showing egg-production on Day 12 following FSC lineage-specific (GAL4 109-30) knockdown of mCherry, Drp1, or Opa1 (N : 5/genotype/condition) L) Bar plot indicating % clonal GSC abundance in CD and HSD (N(germaria)-Stimulated-CD: WT- 9, *drp1^2^*-14, *drp1^1^*-11, *drp1^KG^*-12; N(germaria)-Stimulated-HSD: WT-11, *drp1^2^*-16, *drp1^1^*-17, *drp1^KG^*-11) M) Dot plots showing clonal GSC abundance in each germarium with 0-, 1-, and 2-GSCs where frequency (%) denoted in parentheses for each genotype; gray triangle depicting decreasing Drp1 activity across the mutants. (N(germaria)-Stimulated-CD: WT-9, *drp1^2^*-14, *drp1^1^*-11, *drp1^KG^*-12; N(germaria)-Stimulated-HSD: WT-11, *drp1^2^*-16, *drp1^1^*-17, *drp1^KG^*-11) N) Boxplots showing % clonal differentiated cysts in each ovariole in CD and HSD (N(ovariole)- Stimulated-CD: WT-8, *drp1^2^*-14, *drp1^1^*-11, *drp1^KG^*-12; N(ovariole)-Stimulated-HSD: WT-8, *drp1^2^*- 15, *drp1^1^*-18, *drp1^KG^*-11) O) Dot plot depicting clonal cyst sizes in CD and HSD (N(cyst)-Stimulated-CD: WT-3, *drp1^2^*-7, *drp1^1^*-14, *drp1^KG^*-10; N(cyst)-Stimulated-HSD: WT-10, *drp1^2^*-17, *drp1^1^*-7, *drp1^KG^*-6) P) Stacked bar plots indicating clonal FSC abundance(black) and Loss(grey) (%) in CD and HSD (N(ovariole)-Stimulated-CD: WT-21, *drp1^2^*-20, *drp1^1^*-14, *drp1^KG^*-18; N(ovariole)-Stimulated-HSD: WT-19, *drp1^2^*-18, *drp1^1^*-20, *drp1^KG^*-17) Q) Scatter plot showing FSC Self-renewal / Proliferation for each genotype in CD and HSD (N(cyst)-Stimulated-CD: WT-102, *drp1^2^*-91, *drp1^1^*-102, *drp1^KG^*-76; N(cyst)-Stimulated-HSD: WT- 86, *drp1^2^*-95, *drp1^1^*-114, *drp1^KG^*-102)

**Supplementary Fig 2.**
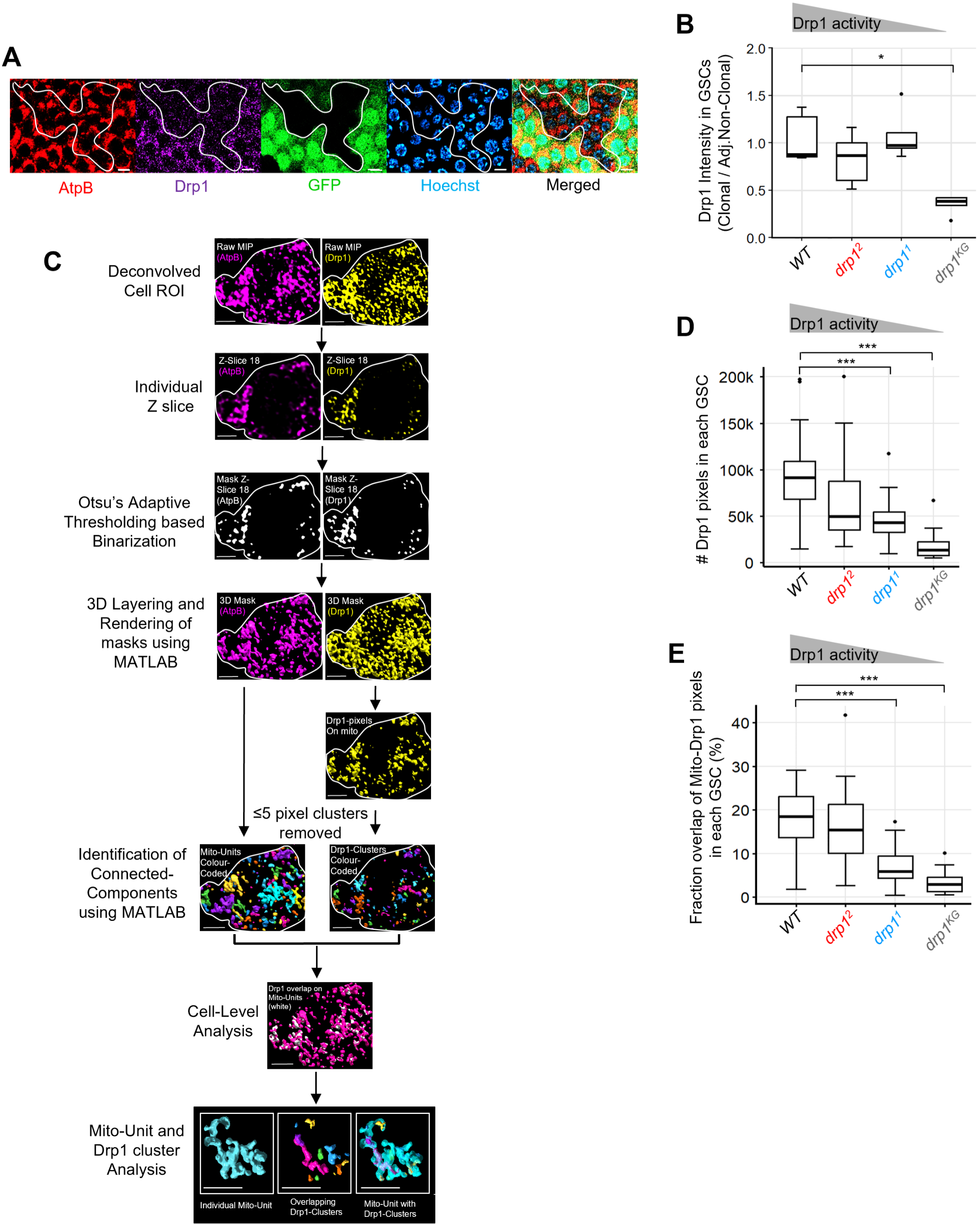
A) Representative micrograph of drp1^KG^ Follicle cells (FCs) stained for AtpB, Drp1 and nuclei. Clonal FCs are highlighted to demonstrate the loss of Drp1 signal within the clonal region compared with the adjacent non-clonal region exhibiting normal Drp1 levels; scale bar : 5 μm. **B)** Box plot showing the ratio of Drp1 intensity between clonal and adjacent non-clonal GSCs for each genotype (N(GSC) : 5/genotype) **C)** Description of the connected-component-based cluster identification pipeline for mitochondrial- units and Drp1-clusters, outlining the sequential steps post image acquisition and preprocessing to extraction of cell-level and cluster-level quantitative parameters; scale bar : 2 μm. **D)** Boxplots showing total Drp1 pixels in each GSC (N(GSC)-Stimulated: WT-21, *drp1^2^*-20, *drp1^1^*- 22, *drp1^KG^*-24) **E)** Boxplots showing the fraction of Mitochondria-Drp1-overlap per mitochondrial-pixels in each GSC (N(GSC)-Stimulated: WT-21, *drp1^2^*-20, *drp1^1^*-22, *drp1^KG^*-24)

**Supplementary Fig 3.**
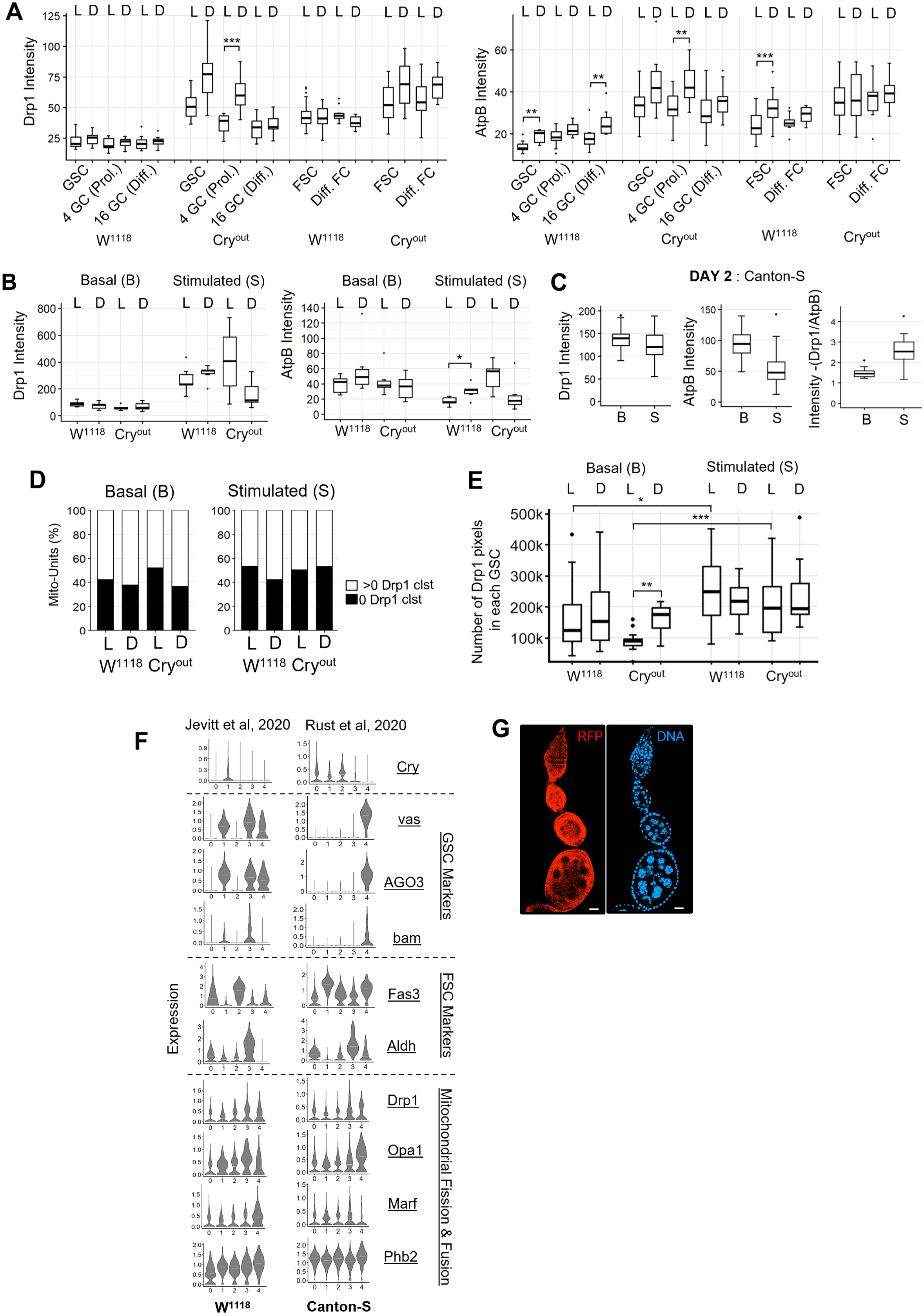
A) Boxplots showing the Drp1 and AtpB intensity in GSC and FSC lineages in the germaria of W^1118^ and Cry^out^ in mid-light(L) and mid-dark(D) in specified developmental stages in basal condition, as quantified from experiment described in 3B. (N-Basal-10/cell type/genotype/time point; except FSCs_N-20) **B)** Boxplots showing the Drp1 and AtpB intensity in GSCs of W^1118^ and Cry^out^ germaria in mid-light(L) and mid-dark(D) in basal and stimulated conditions. (N-6/genotype/timepoint/diet) **C)** Boxplots showing the Drp1 and AtpB intensity and Drp1 intensity normalized by AtpB intensity in GSCs of CS in Basal and Stimulated conditions (N(germaria) : Basal - 34; Stimulated-27) **D)** Stacked-barplots of mitochondrial-units (%) with or without Drp1 clusters (N(Miochondrial- units)-Basal: W^1118^- L-610 ,D-709; Cry^out^- L-721, D-587; N(Miochondrial-units)-Stimulated : W^1118^- L-939, D-924; Cry^out^- L-627 , D-988) **E)** Boxplots showing total Drp1 pixels in each GSC (N(GSC)-12/genotype/timepoint/diet) **F)** Violin plots of all mitochondrial-proteins, GSC and FSC markers and Cry in W^1118^ and CS ovaries from sc-RNA Seq datasets of Jevitt et al. 2020 and Rust et al. 2020. **G)** Representative confocal optical slice of Cry-GAL4 and UAS-RFP ovariole showing RFP expression with Hoechst DNA stain; scale bar : 10 μm.

**Supplementary Fig 4.**
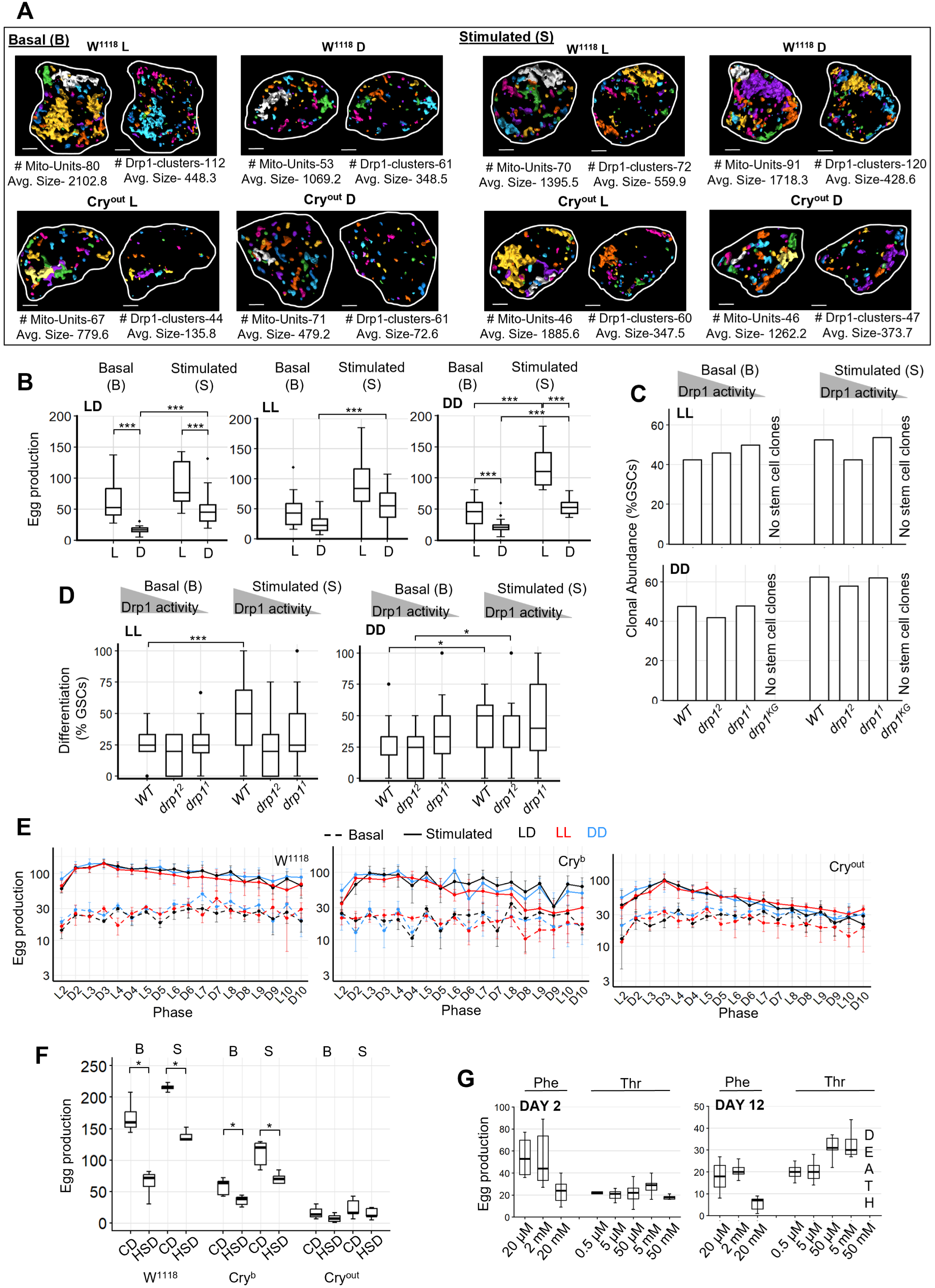
A) Representative section of 3D rendered micrograph of mitochondrial-units and mitochondrially- recruited Drp1-clusters in W1118 and Cryout GSCs mid-light(L) and mid-dark(D) in basal and stimulated conditions. B) Boxplots showing egg-production on Day 12 in CS in basal and stimulated conditions in LD, LL and DD regimes. (N: 15/diet/light regime) C) Bar plot indicating % clonal GSC abundance in LL and DD regimes. Stem cell clones were undetected in *drp1^KG^* (N(GSC)-Basal: LL - *WT*-22, *drp1^2^*-25, *drp1^1^*-24; DD - WT-20, *drp1^2^*-25, *drp1^1^*-25; N(GSC)-Stimulated: LL *WT*-20, *drp1^2^*-25, *drp1^1^*-29; DD- *WT*-19, *drp1^2^*-27, *drp1^1^*- 27) D) Boxplot showing % clonal differentiated cysts in each ovariole. (N(Cyst)-Basal: LL- *WT*-22, *drp1^2^*-25, *drp1^1^*-24; DD- WT-20, *drp1^2^*-25, *drp1^1^*-25; N(GSC)-Stimulated: LL- *WT*-20, *drp1^2^*- 25, *drp1^1^*-29; DD- *WT*-19, *drp1^2^*-27, *drp1^1^*-27) E) Line plot showing egg production in L and D phases in W^1118^ ,Cry^b^ and Cry^out^ from Day 2 to Day 10 in Basal and Stimulated conditions in LD, LL and DD regimes (N: 10 for each diet in each light regime) F) Boxplots showing egg-production on Day 12 in W^1118^,Cry^b^ and Cry^out^ in CD and HSD in basal and stimulated conditions (N: 10 for each diet for each genotype) G) Boxplots showing egg-production on Day 2 and Day 12 in W^1118^ supplemented with Phenylalanine and Threonine (N: 5/Amino acid conc.)

